# Mechanical Force Imprints Neutrophils to Orchestrate Pulmonary Homeostasis

**DOI:** 10.1101/2023.07.03.547316

**Authors:** Jin Wang, Haixia Kang, Wenying Zhao, Wenjuan Bai, Xin Qi, Kepeng Yan, Naijun Miao, Dong Dong, Yuanyuan Wang, Ajitha Thanabalasuriar, Dachuan Zhang, Youqiong Ye, Tian-le Xu, Bin Li, Jing Wang

## Abstract

Neutrophils are the most abundant immune cells that constantly patrol and marginate into the vascular beds of multiple tissues to support immune homeostasis. The extent to which neutrophils undergo adaptation in response to diverse tissue microenvironments, and the resultant biological implications of such adaptation, remains unclear. Here, we performed intravital imaging, transcriptional, and functional analyses of neutrophils in different tissues. Our findings showed that the lung harbors a transcriptionally distinct neutrophil population with unique migratory behaviors. These resident-like neutrophils guarantee a rapid response upon infection and play an important role in maintaining the homeostasis of pulmonary vasculature. Notably, pulmonary neutrophils are imprinted by mechanical cues via the mechanosensitive ion channel Piezo1. Mice with conditional Piezo1 ablation lost lung-specific neutrophil signatures, and showed impaired lung capillary angiogenesis. Furthermore, these mice displayed increased susceptibility to airway infection. Collectively, these data identify mechanical sensing via Piezo1 as an important driver for tissue-specialized neutrophils that support pulmonary homeostasis.

## Introduction

Neutrophils constitute a significant proportion of circulating leukocytes in both humans and mice, accounting for approximately 50-70% and 10-25% of the total leukocyte count, respectively. Known as terminally differentiated and short-lived cells in peripheral blood, neutrophils have been suggested to have limited plasticity and heterogeneity. However, a growing body of research using deep profiling techniques has suggested that distinct neutrophil subpopulations exist in steady state, infectious and inflammatory diseases, tissue repair, and cancer (*1*). The drivers that generate these neutrophil subpopulations has not been well characterized.

Many tissue-resident immune cells, including macrophages and innate lymphoid cells, reside in distinct niches across organs, contributing to tissue homeostasis and the timely response to local microenvironment perturbations (*2*). Traditionally, neutrophils are not defined as tissue-resident cells because they continuously circulate in the blood. Nevertheless, under homeostasis, neutrophils can be found in direct contact with endothelial cells (ECs) of certain vascular beds and are retained in certain tissues for a prolonged time. This population of neutrophils is called the “marginated pool”. The tissues that contain these marginated pools are the lung, liver and spleen (*3*). It has been documented that neutrophils can adapt to the tissue environment they infiltrate and acquire specialized phenotypes and functions (*4*). However, whether and how marginated neutrophils receive ’input’ from the tissue microenvironment and the physiological environment cues that shape neutrophil phenotype and functions are not well studied.

The tissue microenvironment consists of various biological, chemical, and physical components that are favorable for optimal organ function. Soluble factors, xenobiotic factors, nutrients, and metabolites have long been recognized as essential environmental cues of the local cellular micro-niches. Conversely, cells are equally sensitive to some physical properties of the environment, including tissue stiffness, physical confinement, and mechanical tension (*5*). Several studies have revealed mechanisms by which cells perceive mechanical cues from the external microenvironment and transduce them into molecular signals and genetic regulations to coordinate the cellular response (*6*). Neutrophils display considerable plasticity in shape and migratory behaviors while patrolling across different organs and experiencing different mechanical environment. However, the contribution of mechanical sensing in regulating neutrophil phenotype and functions remains unclear.

Herein, we performed intravital imaging and integrative behavior analysis of neutrophils from multiple tissues in physiological states. Through this analysis we observed unique migratory behavior and intracellular signaling in pulmonary neutrophils. Further transcriptomics analysis revealed that neutrophils adapt to the local microenvironment and aquire lung-specific features. Additionally, we identified that mechanical sensing via mechanical sensitive ion channel piezo1 drove the phenotypic and functional specialization of pulmonary neutrophils. Moreover, we revealed that neutrophil deficiency of Piezo1 led to the failure of mounting an effective anti-bacterial response and resulted in decreased pulmonary vascularization at an early age. Consequently, we uncovered a tissue adaptation and specialization process for pulmonary neutrophils that is driven by mechanical force.

## Results

### Neutrophil behavioral landscapes in different tissues

Neutrophils encounter changes in physical conditions while traveling within the circulation through different organs. These include changes in vessel diameters, blood flow velocity, and vessel curvatures. Herein, we hypothesized that neutrophils adapt to these changes in physical cues by altering their migration behavior and morphology, which may also reflect an alteration in genetic and protein content. To verify our hypothesis, we performed intravital imaging with Ly6G^tdTom^ transgenic mice, wherein only neutrophils are labeled with tdTomato **(Figs. S1A–B)**. Subsequently, we imaged multiple tissues at a defined time interval and generated a single overlaying frame of 600-s time-stacks **(120 frames) (Fig. S1C)**. Consequently, we found more neutrophils in the liver, lung, and spleen than in other organs, indicating that neutrophils in these organs migrate at low velocity or stay in the tissue. Cells that appeared in only one frame (< 5 s duration), which we named “flash” cells, showed different frequencies in different organs and were excluded from further analysis (**Fig. S1D) (Mov. S1**). Particularly, we focused on the lung, liver, and spleen as these peripheral tissues contain a large number of neutrophils even in homeostatic states, which was further determined by flow cytometry **(Fig. 1A)**. After data processing and quality control, we have tracked in total ∼2500 neutrophils from lung, liver, and spleen. To gain more insight into the behavior and morphological adaptation of neutrophils upon migration into different tissues, we incorporated a recently published ’behavior landscape’ approach to integrating multiple parameters to reflecting cellular dynamics and morphology (*7*) **(Fig. 1B)**. A total of 38 parameters describing features of kinetics and morphology of hundreds of cells from each tissue were extracted for further analysis **(Table S1)**. UMAP revealed distinct populations related to neutrophil dynamics and morphology even at a louvain resolution setting = 0.2. Moreover, these subpopulations can be further divided into subclusters in each tissue under higher resolution settings **(Figs. 1C and S2A)**.

**Fig. 1.**
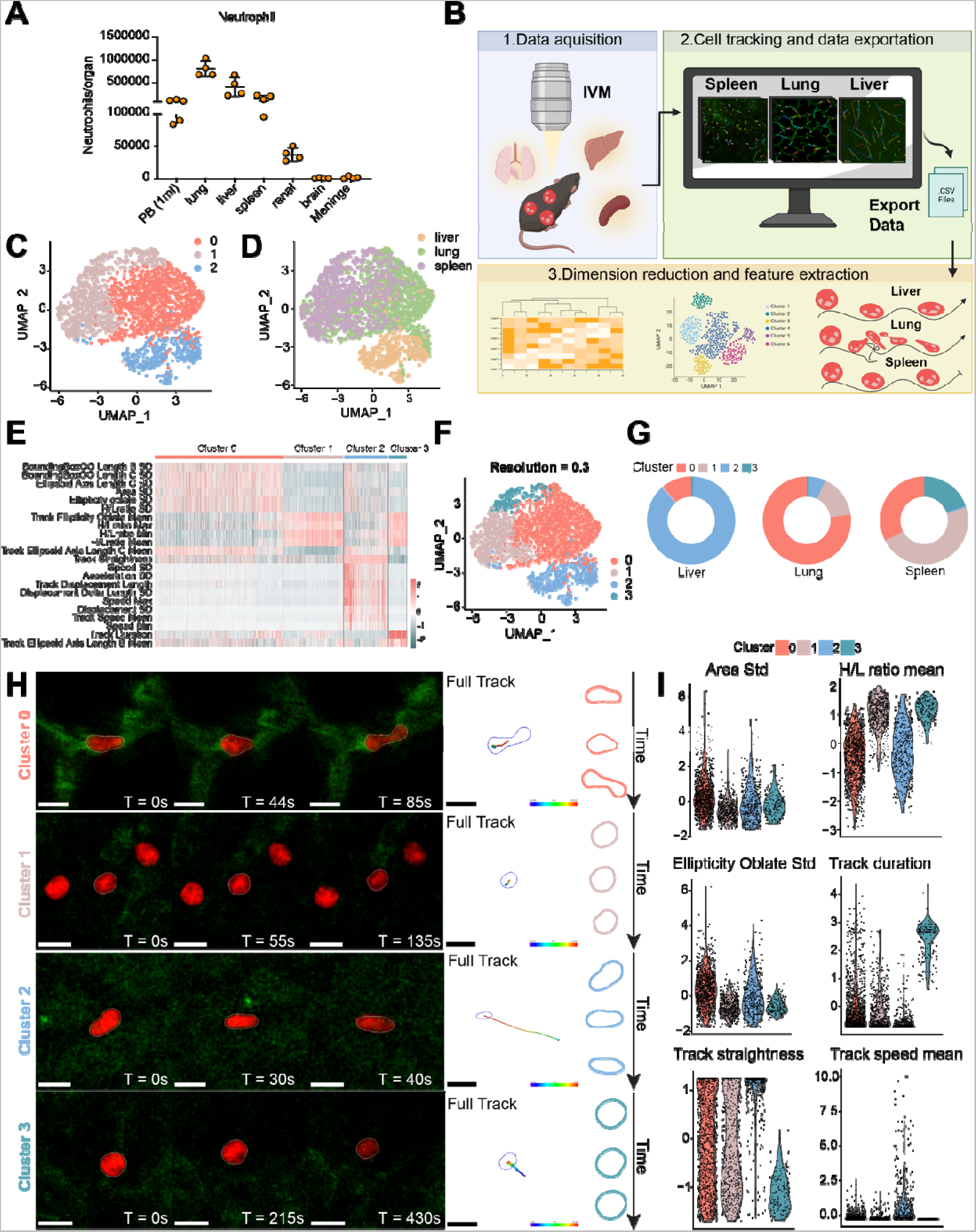
Neutrophil behavior landscapes in different tissues. (A) Neutrophil counts in different tissues. n = 4. (B) Schematic of experiment workflow. Parameters that describe the motion and shape of hundreds of cells were extracted from the analysis of intravital imaging data. A non-linear reduction technique (UMAP) was performed to visualize the data in a low-dimensional space. (C–D) UMAP of 2500 neutrophils from the liver, lung, and spleen. Each dot corresponds to a single cell, colored by cell type (C) or sample origin (D). (E) Heatmap showing the morphology and kinetics parameters for different neutrophil subsets. (F) UMAP of neutrophil subsets, colored according to the clusters identified in (E). (G) Proportions of the four neutrophil clusters in three tissues. (H) Examples of neutrophils from each cluster and their trajectory and shapes at the indicated time points. (I) Violin plots for the indicated parameters across the four behavioral groups, each dot corresponds to a single cell. Data were analyzed using a univariate multinomial model.

Neutrophils within different tissues were then projected into different clusters **(Fig. 1D)**. Subsequently, a resolution setting of 0.3 was determined for the subsequent analysis, as it allows for clear differentiation of neutrophil subsets based on parameters related to migration dynamics and morphology **(Figs. 1E–F)**. The proportion of these subsets in different organs showed consistency between different mice, suggesting the conserved nature of these behavioral traits **(Fig. S2B)**. The major behavioral traits in each tissue are different, implicating an adaptation of neutrophils to different tissue microenvironment **(Fig. 1G)**. These quantitative analyses are largely consistent with our observations. For example, in the lung, 75% of neutrophils showed a prolonged transition time **(Figs. 1H–I and S2C, cluster 0)**. The migration trajectories of these neutrophils are tortuous, and many neutrophils show dynamic deformation (Area Std., H/L ratio mean, and Ellipticity Oblate Std.), which is consistent with the fact that the diameters of lung capillaries are smaller than 5 μm and neutrophils have to change their shape to squeeze through the capillary constriction. About 80% of neutrophils in the liver were migrating with a short transition time and had a more straightforward trajectory **(Fig. S2C, cluster 2)**, while a minor subset showed deformation. The liver sinusoids have an average diameter of ∼10 μm, similar to the diameter of neutrophils. Therefore, most neutrophils flow through the sinusoids without making a stop. However, the liver is known as an organ to clear aged neutrophils through phagocytosis by Kupffer cells lining the vessels (*8*). It is postulated that neutrophils exhibiting slow motility and morphological abnormalities may be the subset that interacts with Kupffer cells. Most splenic neutrophils flow through the blood vessel without deformation **(Fig. S2C, cluster 1)**. A small portion of neutrophils also resides within the parenchyma with static behavior and almost spherical cellular morphology **(Fig. S2C, cluster 3)**. Taken together, we showed that neutrophils could rapidly adapt to different tissue structures and change their behavioral and morphological properties in physiological states. We specifically investigated pulmonary neutrophils, as their abundance and prolonged transition duration suggest that the lung may serve as an unique niche for these cells.

### Neutrophil intracellular calcium signaling in different tissues

Calcium (Ca^2+^) signaling has long been associated with multiple key immunological events, including chemotaxis, phagocytosis, and activation (*9*). The different migratory behaviors might reflect differences in calcium dynamics during neutrophil migration in different tissues. To visualize the calcium event, we expressed the genetically encoded Ca^2+^ indicator Salsa6f (a fusion protein in which tdTomato is linked to the Ca^2+^ indicator GCaMP6f) specifically in neutrophils (Ly6G^Salsa6f^) **(Fig. 2A)**. We used GCaMP6f/tdTomato (G/R) ratio to determine the cytosolic Ca^2+^ levels independent of probe concentration or cell movement. Subsequently, we performed simultaneous calcium, kinetics, and morphology imaging of neutrophils in various tissues. Calcium (Ca^2+^) signalling was detected in neutrophils within the lung, characterized by an augmentation in the intensity of green fluorescence (GCaMP6f) overlaid on red fluorescence (TdTomato). This resulted in transient changes in the apparent color from red to orange and yellow in the whole cell **(Fig. 2B, Mov. S2)**. Subsequently, we performed the ratiometric analysis of GCaMP6f over tdTomato to subtract out fluctuations in green fluorescence due to cell movement. The representative analysis revealed that neutrophils exhibited frequent, short Ca^2+^ transients **(Fig. 2C) (<15 s)**, whereas there are neutrophils in the observed duration exhibited no Ca^2+^ spikes **(Fig. 2D**). Compared to the lung, Ca^2+^ signaling events of neutrophils in liver and spleen are less frequent **(Fig. 2E, Mov. S3)**. In the spleen, most stationary neutrophils remained spherical and did not show calcium signals over time **(Mov. S3)**. Calcium signaling is prevalent in neutrophils during their migration to the lung capillaries but not in the liver, suggesting Ca2+ signaling observed in the lung is unlikely to result from cell migration **(Mov. S3**). We performed a correlation analysis of kinetics and morphological characteristics with Ca^2+^ signal intensities. Neutrophils that were less spherical showed stronger Ca^2+^ signals **(Figs. 2E and S3A)**, whereas there was no direct correlation between Ca^2+^ signal intensity and cell speed, volume, or other parameters **(Fig. S3B)**. The above results suggest that neutrophils exhibit distinct intracellular signaling events in the lung. Spontaneous and evoked Ca^2+^ transients regulate gene expression and functions in neutrophils (*9*). Therefore, we speculate that neutrophils might display a distinct identity and transcriptional programs in the lung.

**Fig. 2.**
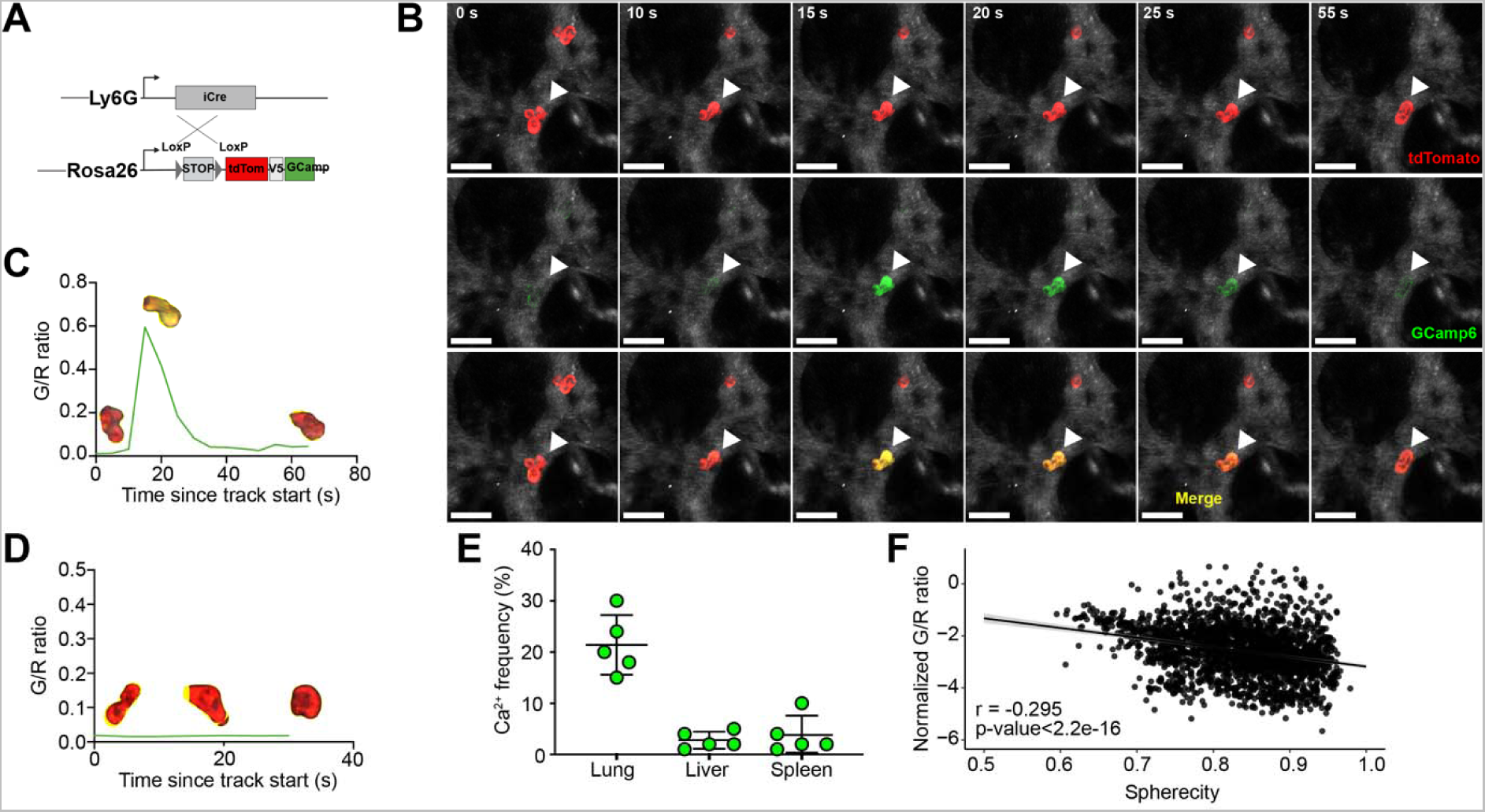
Intravital imaging of Ca^2+^ flux in neutrophils revealed transient Ca^2+^ influx during migration in the lung. (A) Construction strategy of neutrophil calcium reporter mice. (B) Time-lapse images showing a single calcium event in neutrophils in the lung. (C) Mean fluorescence intensity time plot of a neutrophil displaying a short Ca transient. Images of the Ca^2+^ signals of cells are transposed into the plots. (D) Mean fluorescence intensity time plot of neutrophil displaying no Ca^2+^ signal. (E) Frequency of Ca^2+^ signals in neutrophils in different tissues. Data were calculated as the percentage of neutrophils displaying Ca^2+^ in total neutrophils during the same recording time. n = 5 mice. (F) Correlation between GCaMP6f/tdTomato intensity ratio (G/R ratio) and sphericity in neutrophils in vivo.

### Pulmonary neutrophils have distinct mRNA and protein profiles

To further explore whether pulmonary neutrophils possess tissue-specific transcriptional programs, we isolated neutrophils from thoroughly perfused lung, peripheral blood, and bone marrow to perform bulk RNA-seq analysis **(Fig. 3A)**. Analysis of the principal component of gene expression showed that neutrophils clustered according to tissue and lung neutrophils were at greater distances from bone marrow neutrophils compared to blood neutrophils **(Fig. 3B)**. This was confirmed by Pearson correlation coefficients **(Fig. S4A)**. Since neutrophils are generated in the bone marrow before being released into circulation, our subsequent analysis focused exclusively on neutrophils obtained from the blood and lungs. Hundreds of differentially expressed genes (DEGs) were observed in each tissue neutrophil **(Figs. 3C–D) (Table S2)**. Gene ontology (GO) analysis was used to identify enriched pathways in different neutrophil populations. Peripheral blood neutrophils were enriched for pathways, including autophagy, definitive hemopoiesis, DNA repair, myeloid cell differentiation, and myeloid cell homeostasis **(Figs. 3E and S4B)**. The pathways with positive enrichment in pulmonary neutrophils were found to be related to bactericidal function, including response to molecules of bacterial origin, the ROS metabolic processes, and regulation of phagocytosis. The pulmonary neutrophils were also enriched in vasculature development pathways **(Figs. 3F and S4C)**. Notably, pulmonary but not peripheral neutrophils express vascular endothelial growth factor A (VEGFa), a well-known pro-angiogenic factor **(Fig. 3D)**. Therefore, these data suggested that neutrophils underwent tissue-specialized transcriptional reprogramming, which may also impact their functions.

**Fig. 3.**
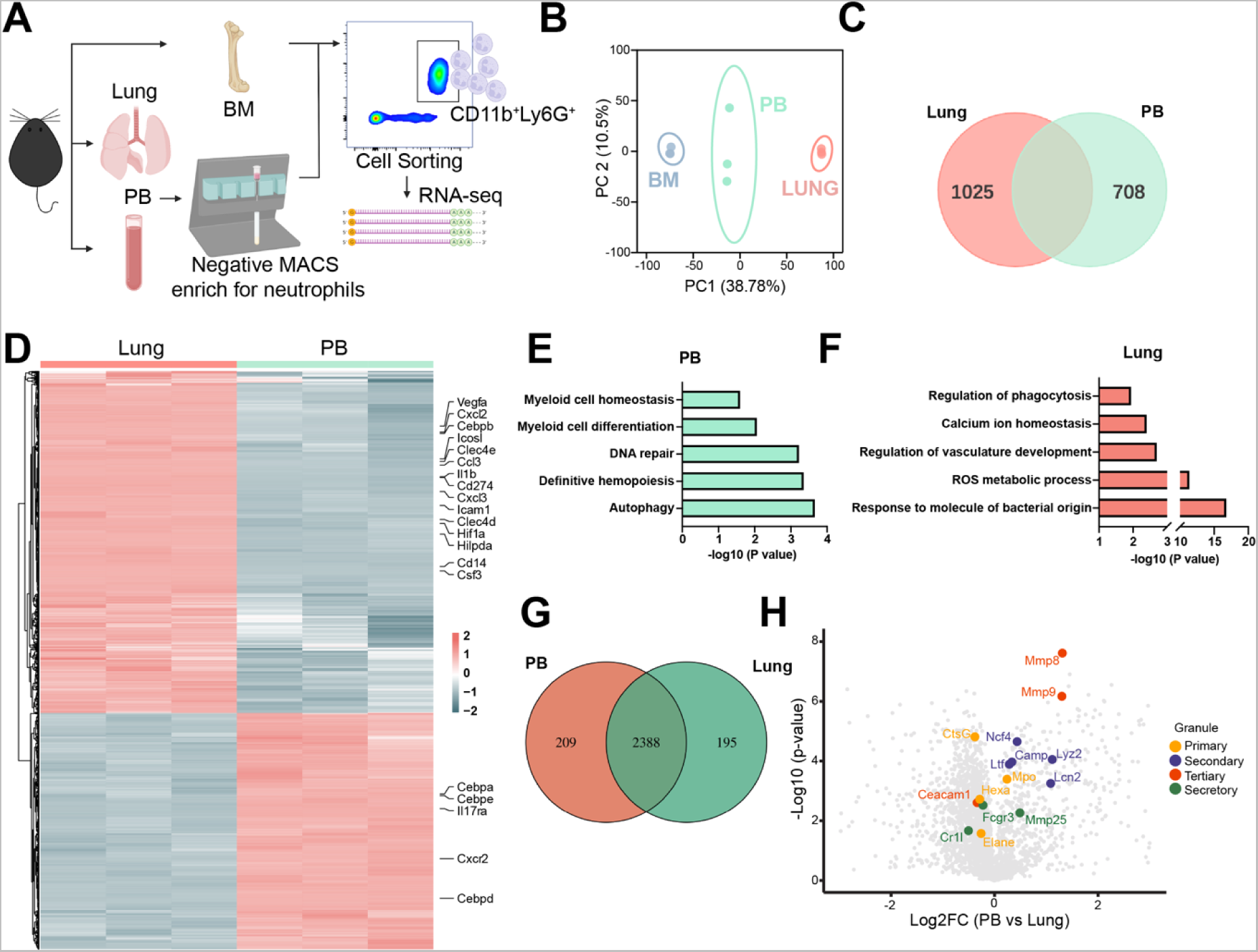
Neutrophils in the lung display distinct transcriptional and proteomic features. (A) Schematic of experiment workflow. (B) Total transcripts of sorted bone marrow (BM), lung, and peripheral blood (PB) neutrophils are subjected to a PCA plot. n = 3 samples per group. (C) Venn diagram representing DEGs between lung and PB neutrophils. (D) Heatmap of DEGs between lung and PB neutrophils, with several representative genes, indicated at right. (E and F) Pathway analysis of DEGs between lung and PB neutrophils. (G) The Venn diagram represents the unique and shared proteins between lung and PB neutrophils. n = 3 samples per group. (H) Volcano plot displaying the distribution of the total proteome from lung and PB neutrophils with relative protein abundance (log_2_FC PB vs. lung) plotted against its significance level (negative log_10_*P*-value), highlighting granule proteins that are significantly (*P* < 0.05) enriched in lung and PB. Colors show the granule type of each protein.

Many proteins are processed and stored for later use during neutrophil development in the bone marrow (*10*). Next, we performed proteomic profiling of blood and lung neutrophils using label-free proteomics. A total of 2792 proteins were identified from blood and pulmonary neutrophils, of which 2693 were common between the two groups **(Fig. 4G) (Table 3)**. The comparison of neutrophil proteome with transcriptional datasets associated with tissue-specific signatures revealed poor correlation **(Figs. S5A–B)**. The proteomic data suggested that pulmonary neutrophils contain more primary granule proteins but fewer secondary and tertiary ones **(Fig. 3H)**. These unbiased and systemic approaches revealed that pulmonary neutrophils expressed unique transcripts and proteins that might equip them for specialized local functions.

**Fig. 4.**
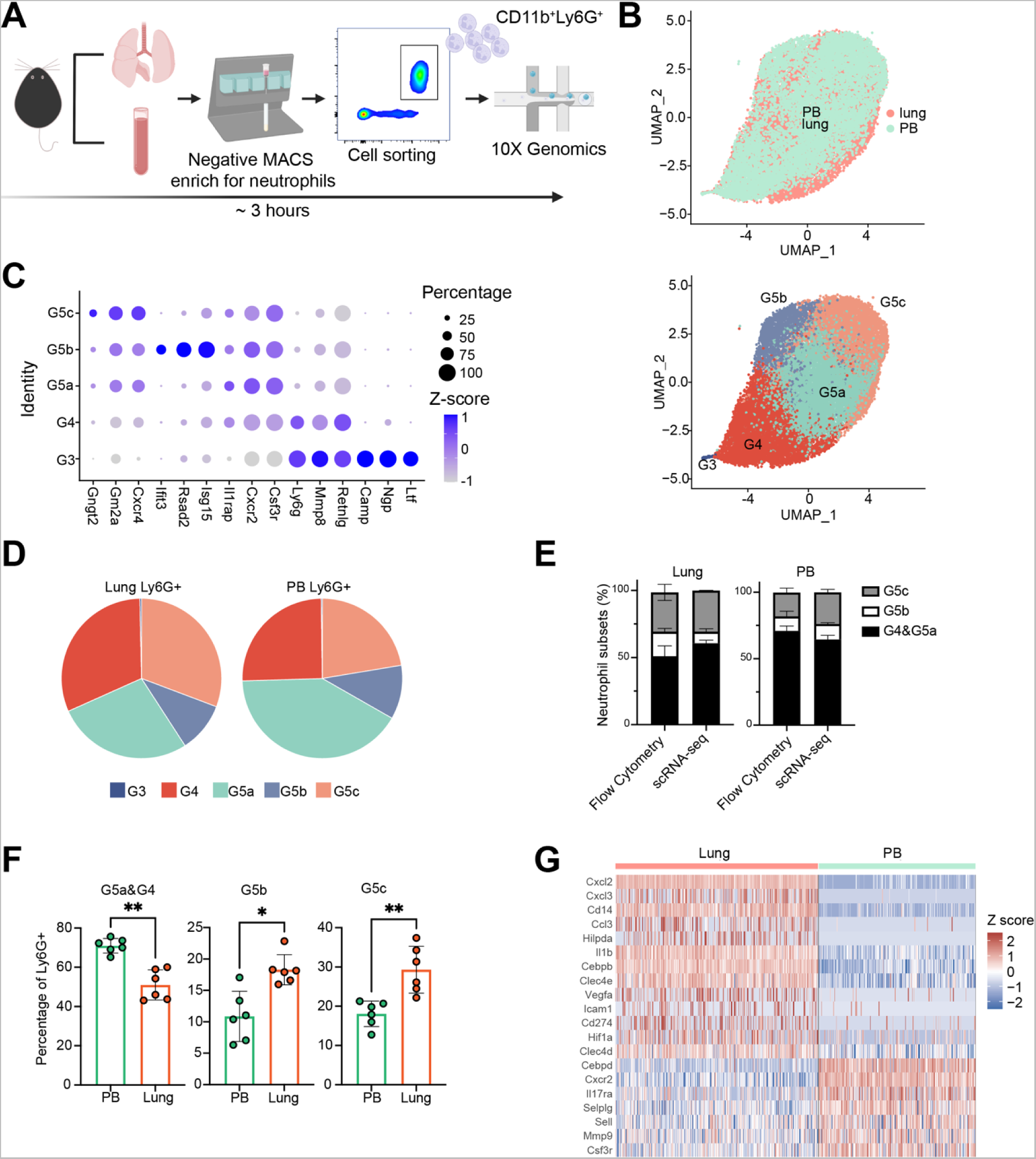
scRNA-seq analysis of neutrophil subpopulations in lung and blood. (A) Schematic of the experimental workflow. (B) UMAP of 20941 neutrophils from PB (9118) and Lung (11823), colored by sample origin (upper panel) or cell type (lower panel). (C) Dot plot showing the scaled expression of selected signature genes for each cluster, colored by the average gene expression in each cluster scaled across all clusters. Dot size represents the percentage of cells in each cluster with more than one read of the corresponding gene. (D) Proportions of the five neutrophil clusters in lung and PB samples. (E) Comparison of FACS and scRNA-seq-based analyses of G4 and G5 subpopulations in PB and lung. (F) Comparison of FACS-based analyses of different neutrophil populations between lung and PB. n = 6 mice. (G) Heatmap showing row-scaled expression of top DEGs from scRNA data for each averaged tissue profile.

### Single-cell profiling reveals conserved neutrophil subpopulations in both the blood and lung

The transcriptional heterogeneity of neutrophils among different tissues prompted us further to explore neutrophil diversity in peripheral blood and lung using single-cell transcriptome profiling **(Fig. 4A)**. After stringent data quality control, we obtained 20941 high-quality cells with an average of 1287 genes per cell profiled **(Fig. S6A) (Table S4)**. Unsupervised clustering identified five major cell populations **(Fig. 4B)**. Subsequently, we compared the cluster identities of our dataset with the scSEQ data in a recent publication (*11*) and found very good agreement in cluster identities and annotations **(Figs. S6B–D)**. Therefore, we used the same annotations as previously reported. We used previously reported signature genes to distinguish each subpopulation and reveal that PMN in circulating blood and lung can be classified into five cell subgroups, including the less mature G3, G4 subsets, and the mature G5a-c subsets. This finding highlights the conserved nature of these neutrophil subpopulations **(Figs. 4B–C) (Table S5)**. However, the proportions of G4 and G5c are higher in the lung compared to blood, whereas G5a is higher in the blood **(Fig. 4D)**. We also calculated the percentage of neutrophil subpopulation in PB and lung using flow cytometry as previously described (*11*) **(Fig. S6E)**. The proportions of G4/G5c, G5b, and G5c in PB and lung measured by FACS analysis are consistent with those calculated based on scRNA sequence data **(Fig. 4E)**. Furthermore, FACS analysis confirmed the higher G5c subset in the lung **(Fig. 4F)**. Although the neutrophil subpopulations were conserved, many genes underwent profound changes in the lung **(Fig. 4G) (Table S6)**. Nonetheless, most of these lung-specific signatures were evenly found in each G5 subpopulation and slightly lower in the G4 subset **(Fig. S7)**. The G4 subset was previously described as newly mature neutrophils in the bone marrow, indicating that the G4 subset may be less mature than the peripheral G5 subsets.

However, despite a quantitative difference among the G4 and G5 subsets, it is unlikely that lung-specific gene signatures are restricted to any subpopulation. Consequently, in the subsequent experiments, we studied lung neutrophils as a whole population. Collectively, single-cell sequencing of peripheral blood and lung neutrophils has confirmed the presence of conserved subpopulations and tissue-specific signatures in lung neutrophils.

### Neutrophils provide immediate protection against airway infection

A previous study has shown that lung neutrophils are an important defense hub for detecting and capturing bloodstream bacteria (*12*). It is unclear how these lung neutrophils respond to airway infection, the main route of infection that causes pneumonia. Mice were challenged with 5 × 10^6^ CFU *Streptococcal pneumonia* (*S. pneumonia*, D39) intratracheally, and the lungs were surgically exposed for intravital imaging **(Fig. 5A)**. As early as 30 min after bacteria inoculation, the pulmonary neutrophils displayed a dramatic behavior change from “jumping” to “crawling” cells **(Mov. S4)**. Using calcium reporter mice revealed continuous fluctuations in calcium across the migrating cells **(Fig. 5B, Mov. S4)**. Calcium signaling frequency and duration of calcium signaling within neutrophils increase significantly, indicating rapid neutrophil activation **(Figs. 5C–D, Mov. S4)**. To further confirm the rapid response of pulmonary neutrophils, we challenged mice with *S. pneumonia* by intratracheal injection and collected bronchial lavage fluid (BALF), lung tissue, and blood at 60 min after infection **(Fig. 5E)**. We observed a significant infiltration of neutrophils in BALF and a concomitant decrease in neutrophil numbers in lung parenchyma after challenge. Conversely, the circulating neutrophils remained the same **(Figs. 5F–H)**. Therefore, these lung ’resident’ neutrophils provide immediate protection against airway infection before circulating neutrophils are recruited.

**Fig. 5.**
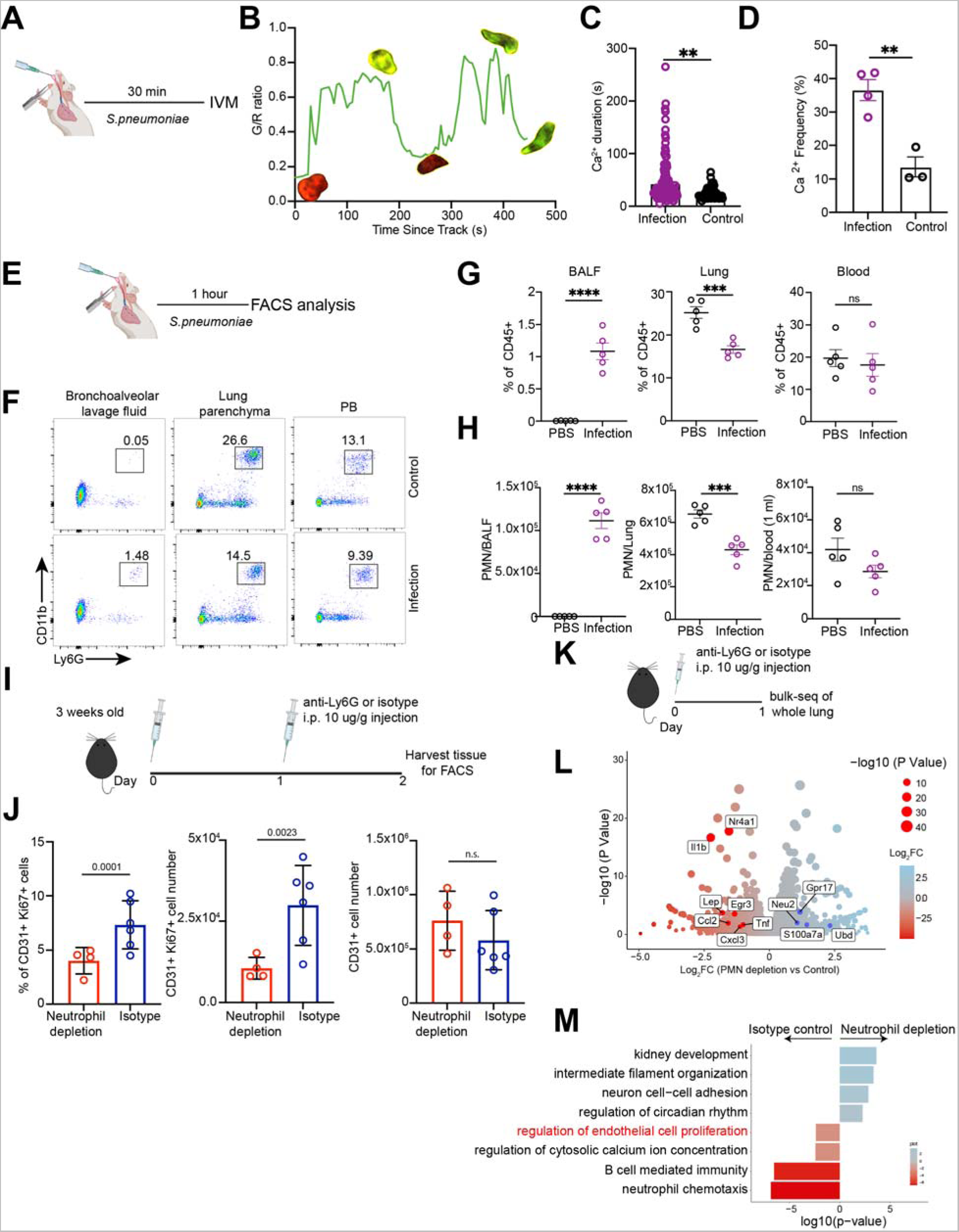
Pulmonary neutrophils are essential for host defense and maintenance of vascular homeostasis. (A) Schematic of the experimental workflow. (B) Representative mean fluorescence intensity time plot of pulmonary neutrophil displaying Ca^2+^ signals following i.t. infection in vivo. Images of the Ca^2+^ signals of cells are transposed onto the plots. (C–D) Duration (C) and frequency (D) of Ca^2+^ signals in neutrophils in PBS (control) and infected lung (infection), as assessed by intravital imaging. n = 4. (E) Schematic of the experimental workflow. (F) Representative flow cytometry analysis of neutrophils in different tissues. Cells were pregated on size, live/dead, and CD45^+^. (G–H) Percentage (G) and cell numbers (H) of neutrophils in different PBS and bacteria-challenged mice tissues. n = 5. (I) Schematic of the experimental workflow. (J) Volcano plot showing DEGs between isotype control and neutrophil-depleted lungs. Transcripts significantly upregulated in PMN depletion and control lungs are colored in blue and red, respectively (log_2_fold change ± 0.5 and adjusted *P* value < 0.05). (K) GO enrichment analysis was performed on the DEGs. Blue indicates enrichment in lungs from neutrophil depletion mice, while red indicates enrichment in lungs from control mice. (L) Schematic of the experiment workflow. (M) Flow cytometry analyzed the percentage and number of CD31+ Ki67+ cells and the total number of CD31+ cells in isotype control and neutrophil-depleted lungs. n = 4 (neutrophil depletion) or 6 (isotype).

### Neutrophils promote pulmonary vascular angiogenesis

An important function previously documented to be specialized for lung neutrophils is their contribution to pulmonary vascularization (*4*). Similar to previous report, neutrophil depletion at an early age resulted in a complete loss of the proliferative capacity of endothelial cells **(Figs. 5I–J)**. To investigate whether neutrophil depletion may have a broader influence in the lung tissue, we performed bulk RNA sequencing of the entire lung tissue in neutrophil-depleted mice **(Fig. 5K)**. Long-term neutrophil depletion has been reported to affect the progress of hematopoiesis and natural killer cell homeostasis (*13, 14*). We only performed short-term neutrophil depletion and harvested tissue at 24 h to avoid the following cascading effects caused by other immune compartments. Antibody treatment efficiently depleted neutrophils without affecting monocytes in the peripheral blood and lung **(Figs. S8A–B)**. The analysis of transcriptomic data revealed that neutrophil depletion affected many genes that could be classified into specific pathways **(Fig. 5L) (Table S7)**. Consistent with efficient neutrophil depletion, genes related to neutrophil chemotaxis (Csf3r, Cxcr2, Cxcl3, Ccl2, Ccl4) were downregulated. Neutrophil depletion has been observed to impact various biological processes such as regulation of endothelial cell proliferation and cytosolic calcium ion and B cell-mediated immunity **(Fig. 5M)**. These findings indicated that neutrophils play an important role in regulating pulmonary vascular homeostasis and may also potentially affect other biological processes in the lung.

### Pulmonary neutrophils respond to mechanical cues via Piezo1

Our imaging data revealed that neutrophils continuously squeeze into numerous small vessels during their transition through the pulmonary circulation and repeatedly experience significant physical strain. The observed phenomenon prompted us to explore whether mechanical sensing by neutrophils might play a role in shaping lung-specialized neutrophil functions. Consequently, we examined the expression levels of known mechanosensitive ion channels in neutrophils. Piezo1, and to a lesser extent, Trpv4 were abundantly expressed in neutrophils **(Fig. 6A)**. The expression of Piezo1 in lung neutrophils was confirmed using Piezo1P1tdT mice, in which a Piezo1-tdTomato fusion protein is expressed **(Fig. 6B)**. The conditional Piezo1-deficient mice were generated by crossing mice expressing Cre recombinase directed by the vav1 promoter with mice expressing homozygous loxP sites around exons 20–23 of Piezo1 (*Piezo1*^fl/fl^); hereafter referred to as *Piezo1*^ΔVav1^. The expression of Piezo1 in neutrophils was significantly ablated at transcriptional and protein levels in *Piezo1*^ΔVav1^mice **(Figs. S9A–B)**. Live cell imaging revealed that PIEZO1 agonist Yoda1 stimulation triggered a transient calcium influx in wild-type (*Piezo1*^fl/fl^) neutrophils but not *Piezo1*^ΔVav1^ neutrophils **(Fig. 6C) (Table 8)**. The whole cell patch clamping recording revealed the neutrophil response to pressure **(Fig. 6D)**, which was completely abolished in the absence of Piezo1 **(Fig. 6E**). To determine whether piezo1-mediated mechanical sensing contributes to neutrophil dynamics in vivo, we performed intravital imaging to compare migratory behaviors, morphology, and Ca^2+^ signaling of pulmonary neutrophils in wild-type and *Piezo1*^ΔVav1^ mice that also express Salsa6f **(Mov. S6)**. In Piezo1 depletion mice, more neutrophils displayed short transient time (defined as “flash” cells) compared to the wild-type control **(Figs. 6F–G)**. Piezo1-deficient neutrophils also displayed a lower Ca^2+^ frequency when migrating in the lung **(Fig. 6H)**. The analysis of duration time distribution of the observed neutrophils revealed a higher frequency of neutrophils with a short duration time in Piezo1-deficient mice. This observation suggests that Piezo1-deficient neutrophils spent a shorter time passing through the lung **(Fig. 6I)**. The changes in these migration features and intracellular signaling events highlight the essential role of mechanical sensing in regulating neutrophil behaviors in the lung. These results demonstrated that neutrophils respond to mechanical cues in the lung via piezo1. Moreover, piezo1 is essential for controlling neutrophil deformation and migration behavior in the lung.

**Fig. 6.**
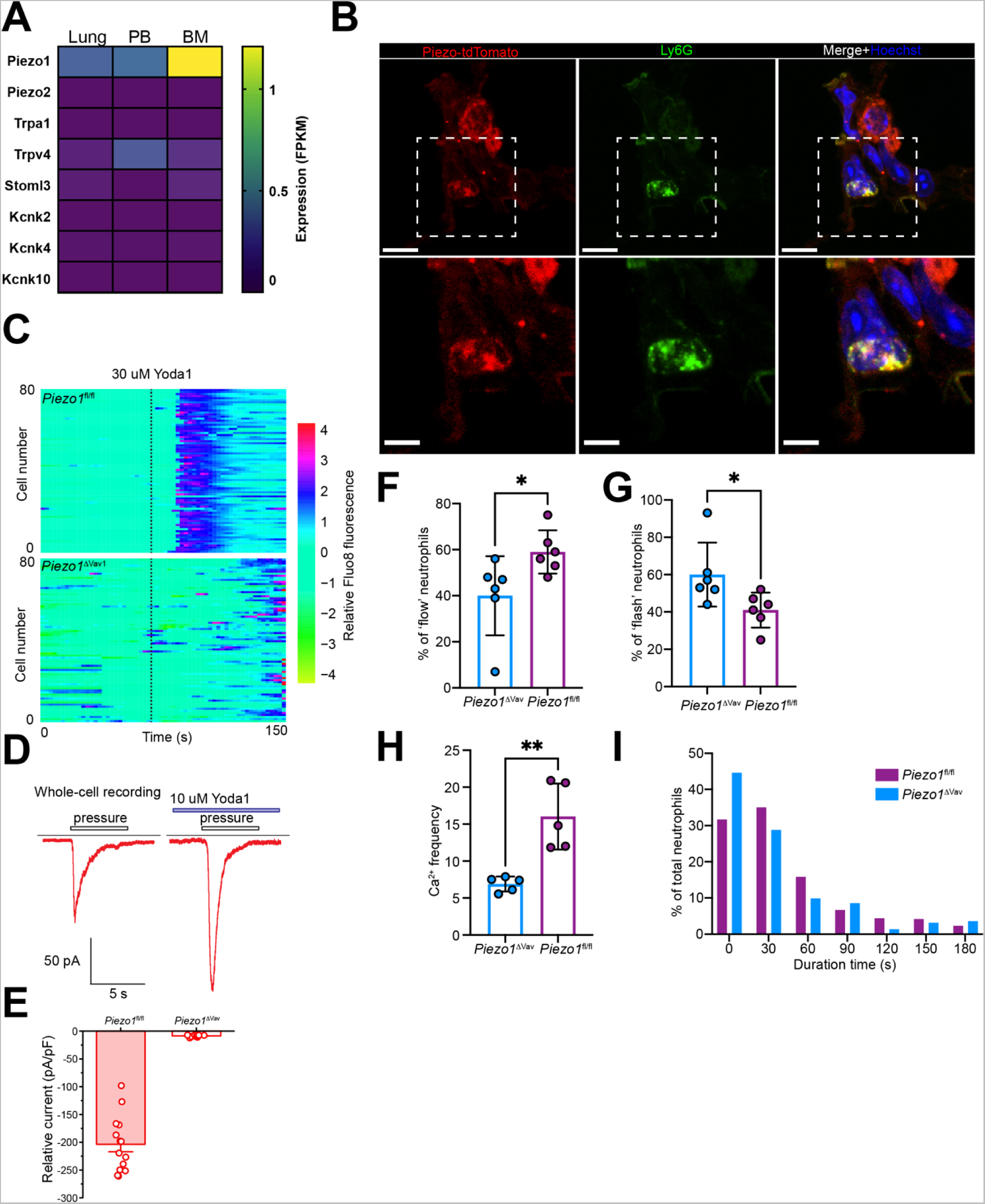
Neutrophils in the lung respond to mechanical cues via Piezo1. (A) Heatmap of different mechanical sensing receptors in neutrophils from different tissues. Data were generated from the log_2_FPKM of the mean value of replicates from RNA-seq data in Fig. 3. (B) Immunostaining of Piezo1-tdTomato and the neutrophil marker Ly6G from lung slice derived from the Piezo1-tdTomato-KI mice. The enlarged view of regions highlighted in the dashed box is shown below. Hoechst was used for staining the nucleus. Scale bars, upper panel: 10 μm; lower panel: 5 μm. (C) Heatmap of normalized calcium fluorescence versus time for each single bone marrow neutrophil isolated from wild-type (*Piezo1*^fl/fl^) and Piezo1 deficient (*Piezo1*^ΔVav1^) mice. (D) Representative whole-cell currents of neutrophils in response to poking pressure and adding Yoda1. (E) Whole-cell currents from neutrophils of *Piezo1*^fl/fl^ and *Piezo1*^ΔVav1^ mice in response to poking pressure. (F–G) Percentage of “flow” and “flash” neutrophils per field of view, as assessed by intravital imaging of lungs from *Piezo1*^fl/fl^ and *Piezo1*^ΔVav1^ mice. Data are from 6 mice per group. (H) Frequency of Ca^2+^ signals in neutrophils, as assessed by intravital imaging of lungs from *Salsa6f*^ΔVav1^ and *Salsa6f*-*Piezo1*^ΔVav1^ (I) Distribution of track duration for lung neutrophils in *Piezo1*^fl/fl^ and *Piezo1*^ΔVav1^ mice. n > 250 tracks per group, three imaging fields of each mouse, each group involves thre mice.

### Mechanical sensing via piezo1 drives lung-specific neutrophil programming

To further explore whether PIEZO1-mediated mechanical sensing drives the lung specialized phenotype and functions in neutrophils, we performed a bulk RNA-seq in lung and blood neutrophils from *Piezo1*^fl/fl^ and *Piezo1*^ΔVav1^ mice **(Fig. 7A)**. The PCA analysis revealed a profound transcriptional difference between *Piezo1*^fl/fl^ and *Piezo1*^ΔVav1^ neutrophils in the lung compared to those in the blood **(Fig. 7B)**. There were only a few overlaps in the DEGs between *Piezo1*^fl/fl^ versus *Piezo1*^ΔVav1^ neutrophils in the blood and lung **(Fig. 7C) (Table S9)**. Notably, DEGs in *Piezo1*^fl/fl^ and *Piezo1*^ΔVav1^ pulmonary neutrophils showed a significant correlation to DEGs between wild-type PB and lung neutrophils **(Fig. 7D) (Table S10)**. The GO analysis revealed the effect of Piezo1 on many biological pathways, many of which were also enriched in lung neutrophils when compared with blood **(Fig. 7E)**. These results suggested that Piezo1 might control the tissue-specialized identities of lung neutrophils. In particular, genes involved in vascular growth and repair (including VEGFa) were only enriched in wild type and not in those of *Piezo1*^ΔVav1^ **(Fig. 7C)**. Consequently, compared to young age wild-type mice, *Piezo1*^ΔVav1^ mice showed impaired proliferation in lung endothelial cells **(Figs. 7F–H)**. To test whether Piezo1 also contributes to the host defense function of lung neutrophils, we infected *Piezo1*^fl/fl^ and *Piezo1*^ΔVav1^ mice with 2 x 10^6^ colony-forming units (CFU) of *S. pneumoniae* D39 intranasally or 1 x 10^7^ CFU intravenously and monitored the survival rate. The *Piezo1*^ΔVav1^ showed a decreased survival exclusively upon intranasal infection, as opposed to intravenous infection **(Figs. 7I–J)**, indicating that mechanosensation via piezo1 mainly affects the anti-bacterial response of pulmonary neutrophils. Collectively, these data demonstrate an important role for PIEZO1 in mediating tissue adaptation of neutrophils in the lung, and this adaptation is essential to maintain pulmonary homeostasis.

**Fig. 7.**
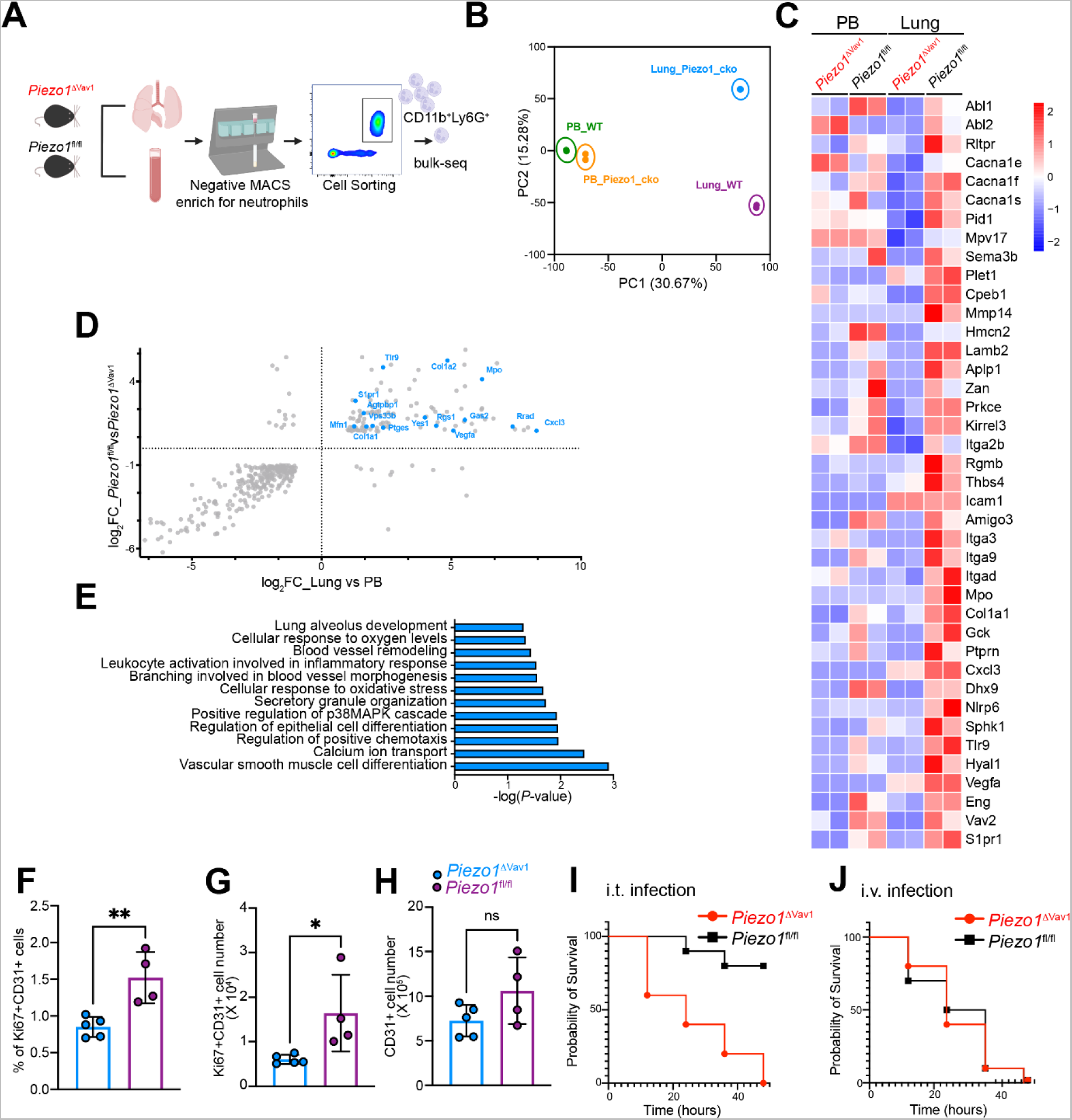
Mechanical sensing reprograms pulmonary neutrophils via Piezo1. (A) Schematic of experimental workflow. (B) PCA of gene expression data of neutrophils isolated from the blood and lung of wild-type (WT) and *Piezo1*^ΔVav1^ (Piezo1_cko) mice. (C) Heatmap of DEGs between *Piezo1*^fl/fl^ and *Piezo1*^ΔVav1^ neutrophils in PB and lung. (D) Correlation of gene expression between RNA sequencing datasets (*Piezo1*^fl/fl^ versus *Piezo1*^ΔVav1^ pulmonary neutrophils, y-axis; lung versus blood neutrophils, x-axis). Only genes with differential expression in both datasets are shown. (E) Pathway analysis of DEGs is highlighted in (D). (F–G) Percentage (F) and absolute count (G) of proliferating endothelial cells in lungs from three weeks old *Piezo1*^fl/fl^ and *Piezo1*^ΔVav1^ mice. (H) Absolute count of total endothelial cells in lung from three weeks old *Piezo1*^fl/fl^ and *Piezo1*^ΔVav1^ mice. (I–J) Survival of *Piezo1*^fl/fl^ and *Piezo1*^ΔVav1^ mice that were infected with D39 via i.t. (I) or i.v. (J) route. n = 10 per group.

## Discussion

Neutrophil heterogeneity and plasticity have recently gained much attention (*15*). Emerging technology, including single-cell sequencing and CyToF, has been used to demonstrate different subsets of neutrophils in mice and humans based on differences in transcriptional and protein levels (*4, 11, 16*). Although some level of heterogeneity between neutrophils from different tissues has been recognized, the concept of tissue adaptation for neutrophils has not been well studied. Neutrophils are characterized by their high degree of dynamism and are exposed to diverse tissue environments while circulating. The majority of neutrophils remain in a state of equilibrium within the bloodstream, with the exception of those located in the spleen, lymph nodes, and intestinal tissue (*17*). It is unlikely that neutrophils will receive as many “imprinting” signals as those for macrophages from the local tissue parenchyma because the blood vessel barrier physically separates them.

Conversely, vascular architecture and hemodynamics might be the most important factors determining neutrophil identity. Herein, we performed high throughput intravital imaging studies to compare neutrophil migration behavior in different organs. Through performing an unbiased multiparameter behavior landscape analysis, we identified different neutrophil behaviors in the lung based on their kinetics and morphology. The unique behavioral features of neutrophils in the lung further encouraged us to characterize the specific transcriptional and translational features of neutrophils in the lungs using bulk RNAseq and scRNA-seq. These complementary assays provided consistent results and further confirmed the specific signatures of neutrophils in the lung. Therefore, there is an intense interest in integrating data from different layers to gain better insight into the heterogeneity and plasticity of neutrophils in the future.

Large numbers of neutrophils in healthy lung tissue suggest that they perform physiological functions and can adapt to the local environment. Our data showed that neutrophils profoundly change their transcriptome profiles from the bone marrow to the blood and lung, indicating that neutrophils undergo tissue adaptation. A previous study suggested that the CXCR4-CXCL12 axis may form instructing regions or niches where neutrophils receive tissue-derived signals, although it is not clear whether neutrophils will lose their lung-specific features in the absence of this axis (*4*). Here, we focus on the unique anatomic structure of lung. The alveoli are the basic working unit for respiratory function in the lung. A single alveolus is surrounded by ∼ 1000 inter-connected short tubular capillary segments arranged in spheroidal mesh (*18*). This unique structure guarantees efficient gas exchange. A neutrophil is estimated to pass through ∼ 40–100 capillary segments during a single transit from an arteriole to a venule (*19*). Due to the size mismatch, in at least half of these segments, neutrophils are required to deform to get through. The phenomenon of neutrophil deformation during transit through the pulmonary vasculature has been observed and documented for several decades. However, a comprehensive investigation into the biological implications of these observed behaviors has yet to be conducted. (*20*). Our study demonstrates the critical role of these changing mechanical properties in shaping neutrophil identities. Additionally, we identified Piezo1 as the mechanical sensor that mediated neutrophil detection. Several families of molecular mechanoreceptors have been reported to transduce mechanical stimulation into ion currents thereby regulating physiological (*21*). Piezo1 has been identified as a component of mechanically activated cation channels in both vertebrates and non-vertebrates (*22*). In addition to Piezo1, another mechanical sensor TRPV4 has been reported to regulate neutrophil activation in acute lung injury (*23*). It is unclear whether TRPV4 also contributes to neutrophil homeostatic regulation. Furthermore, most neutrophils are in the circulation that is constantly exposed to shear stress, another mechanical input. In the future, it will be intriguing to investigate how different mechanical inputs might affect the neutrophil response.

The function of piezo1 in innate immune cells has recently been documented (*24*). Activation of Piezo1 in monocytes has been related to increased pro-inflammatory gene expression in lung infection, sepsis, pulmonary fibrosis, and renal fibrosis (*25, 26*). Piezo1-mediated silencing of the retinoblastoma gene Rb1 drives immunosuppressive myelopoiesis in cancer and infectious disease (*27*). Elevated Piezo1 activity in neutrophils has been shown to have pathological effects in patients with type 2 diabetes (T2D). Piezo1 activation of Piezo1 in neutrophils from T2D patients triggered the release of extracellular neutrophil traps, which could contribute to neutrophil-driven thrombotic risk (*28*). Our study suggested that physiological mechanical signals imprint pulmonary-specific signatures in neutrophils via piezo1. The lack of piezo1 in hematopoietic cells resulted in impaired capillary angiogenesis at an early age and failed clearance of bacterial infection. It seems that fine-tuning mechanical signaling through piezo1 is required to maintain important immune functions and to avoid detrimental responses. Investigating the threshold force to activate piezo1 under different conditions and the crosstalk between mechanotransduction and other signal pathways will be of interest.

Collectively, we showed that neutrophils display unique transcriptional, protein, and behavioral signatures in the lung. Neutrophils that are trapped in the nearby environment are poised to mount a prompt response to any insult and ensure expeditious clearance. Moreover, pulmonary neutrophils aid in vascular development at an early age. These specialized, enhanced functions of neutrophils are instructed by local mechanical signals, which are generated when neutrophils migrate through the small capillaries. Neutrophils face much resistance when migrating in interstitial tissue during the inflammatory response, where they constantly deform. The contribution of mechanical sensing to the inflammatory response should be a topic of great interest for future investigation.

## Materials and Methods

### Mouse strains

Ly6g-Cre-2A-tdTomato was purchased from Shanghai Model Organisms, whereas Piezo1-tdTomato (029214) and Salsa6f (031968) were purchased from Jackson Laboratory. The Piezo1 fl/fl (0292143) and Vav1-iCre (008610) were generously provided by Dr. Jianwei Wang from Tsinghua University and Dr. Zhaoyuan Liu from Shanghai Jiao Tong University School of Medicine, respectively. The C57/BL6 mice involved in this study were all purchased from SLAC ANIMAL, Shanghai. The mice used in the study were 7–12 weeks-old females or males (no sex difference was observed). The mice were housed under specific pathogen-free conditions at ∼22 with humidity of 40%–70% and a light/dark cycle of 12 h. It is generally recognized that neutrophils have intrinsic circadian oscillations and that they manifest different phenotypes at varying times. As a result, animal experiments typically start in the morning (around 10:00) except for intravital imaging.

### Intravital imaging and data processing

All procedures were performed on mice anesthetized with 75 mg/kg Zoletil^TM^ (Virbac) combined with 18 mg/kg Xylazine hydrochloride (Sigma, X1251). The body temperature was maintained throughout surgery and imaging by placing mice on a heating pad at 37 °C. The mice were injected intravenously with 10 µL Qtracker705 vascular tracker (Invitrogen, Q21061MP) or 30 µL 1% FITC-Dextran70 (TdB Labs, 21024) to visualize vasculature and 7 µL APC-Ly6G (Biolegend, 127614) antibody to label neutrophil 30 min before imaging.

#### Lung

The mice were taped to a custom microscope stage, and a small tracheal cannula was inserted and fixed with suture and attached to a mouse ventilator (RWD, R405). The mouse was ventilated with a tidal volume of 10–12 µL of compressed air per gram of mouse weight, a respiratory rate of 130 breaths per minute, and a positive end expiration pressure of 3 cm H_2_O. After cannulation, the mice were placed in the right lateral decubitus position, and a small surgical incision was made to expose the rib cage. A second incision was made by removing three ribs to expose the left lung lobe. A flanged 3D-printing thoracal window with 4 mm coverslip was placed above the exposed lung surface, and 20 mmHg of suction was applied (Yuyan Instrument) to gently immobilize the lung.

#### Spleen and kidney

To expose the spleen and liver, mice were placed in a right lateral position, and a skin incision was made on the left flank. The same window used for lung imaging was used to facilitate imaging of the spleen and kidney.

#### Liver

A midline incision was performed, followed by a lateral incision along the costal margin to the midaxillary line to expose the liver. Mice were placed in a right lateral position, and the ligaments that connect the liver to the diaphragm and the stomach were cut, allowing the liver to externalize onto a glass coverslip. Exposed abdominal tissues were covered with saline-soaked gauze, and the liver was draped with saline-soaked gauze to prevent dehydration. Finally, the mice were placed on a plate with a coverslip for imaging.

#### Brain and Meninge

The fur on the head and neck was shaved, and the skin on the parietal skull bone was surgically removed. The membrane above the skull bone was carefully removed, and a homemade iron imaging plate was glued to the skull bone using dental cement. The skull bone over the barrel cortex was thinned using an electrical micro-drill (RWD, 800-003777-00) to a thickness of ∼30– 40 um. Second-harmonic signals (SHG) are used to identify meninge, and we used the shape and size of vasculature to discriminate pia mater from the parenchyma.

Time-lapse images were acquired with an upright microscope (OLYMPUS FLUOVIEW FVMPE-RS equipped with an XLPlan N 25×/1.05 objective (Olympus) for lung, spleen, kidney, brain, and meninge) or an invert microscope (OLYMPUS FLUOVIEW VS3000 equipped with a UplanSApo 20x/0.75 objective (Olympus) for liver). Acquisitions were performed at the following excitation wavelengths: 920 nm for FITC and tdTomato, 1150 nm for APC, and 800 nm for SHG. The images were acquired with the following settings: 640 x 640 pixels every 5 s for 10 min. To exclude artificial results (autofluorescence and potential tissue damage caused by surgery), we acquired a single z at least 15 µm below the tissue surface, and all videos were acquired within 1 h after surgery for each tissue. For behavioral landscape, we acquired all videos with a single z; for in vivo calcium imaging, a full z stack for 9 µm at a 3- µm interval every 5 s was captured to generate a 4D image.

### Intravital Ca^2+^ imaging

*Ly6g-Salsa6f*, *Vav1-Salsa6f-Piezo1^-/-,^* and *Vav1-Salsa6f* were used to measure intracellular Ca2 + levels *in vivo* intracellular Ca^2+^ levels. GCaMP6f/TdTomato (green/red (G/R) ratio) was obtained to generate additional traces showing single cell and average changes in cytosolic Ca^2+^ over time. For in vivo calcium imaging, we generated a new channel specifically for the tracked cells by the ’Coloc’ function of Imaris to avoid the influence of lung autofluorescence. Average intensities in the green and red channels were calculated for each tracked neutrophil at each time point and exported as a data matrix. Further analysis and visualization were performed in R programming language. The heat map was plotted with the package pheatmap, whereas the scatter plot and correlation maps were generated by corrplot and ggplot2 packages, respectively. Furthermore, a Spearman rank-order correlation analysis was performed and plotted in R.

### Image Analysis

Image data were analyzed with FIJI and Bitplane Imaris (9.9.1). Imaris was used to detect cells and generate tracks with its built-in algorithm. Furthermore, automated generated tracks were reviewed and corrected manually. Subsequently, we added an area filter to filter cells less than 75 µm^2^.

### Behavioral Landscape

For the behavioral landscape analysis, 118 parameters calculated by Imaris for each track were exported from Imaris and further selected and analyzed in R (3.6.0) using the Seurat (3.2.1) package. Some description parameters such as min, max, and standard deviation were calculated in R through its base function, such as lapply and aggregate function. According to a published work, the input matrix consists of 38 selected parameters and 2584 tracks (1296 from the lung, 867 from the spleen, and 421 from the liver). We performed principal component analysis (PCA), and cells were clustered based on k-nearest neighbor graphs using the Louvain algorithm. Finally, a uniform manifold approximation and projection (UMAP) was performed to visualize the data in a low-dimensional space and displayed through the Seurat built-in function Dimplot. To identify the parameters differentially scored between the two clusters, we selected those that showed at least a 0.25-fold difference (log scale) between the two groups. We used the Seurat built-in functions vlnPlot and Doheatmap to visualize each subset and plot the parameter value distribution for each group.

### Tissue Dissociation

Mice were anesthetized and transcardially perfused with 10 mL of cold PBS. Liver, lung, kidney, and brain were cut into small pieces (< 2 mm^2^) and incubated in 3 mL digestion solution (0.5 mg/mL Collagenase IV and 50 units/mL DNaseI in RPMI-1640 medium supplemented with 10% FBS) for 40 min at 37 °C with 5% CO_2_. To obtain a single-cell suspension, digested tissues were disrupted by mincing through a 70- µm cell strainer, centrifugated, and resuspended with 1 mL of FACS buffer (2% FBS with 1 mM EDTA). The brain-cell suspension was pipetted with a Pasteur tube ten times and was separated by a single layer 37% Percoll gradient and centrifugation at 1500 xg for 25 min at 25 °C with low acceleration and no brake. Cells at the bottom were collected. The dura mater was peeled from the top of skull with fine tweezers and cut into small pieces, incubated in 1 mL of digestion solution with gentle shaking every 10 min, followed by mincing through a 70-µm cell strainer. The spleen was mechanically dissociated using a dounce homogenizer (Sigma, D8938). The cells were filtered through a 70- µm cell strainer, followed by red blood cell lysis via 1 mL ACK (Gibco, A10492) for 2 min at room temperature. BM was collected from the tibias and femurs of both hind legs of the mice. The end of both bones was cut, then centrifugated at 8000 rpm for 1 min to collect BM cells. The pelleted BM cells were resuspended with 200 µL ACK red blood cell lysis solution to remove erythrocytes and incubated at room temperature for 1 min. Total peripheral blood was collected at a volume of 1 mL per mouse before perfusion via the retroorbital sinus with an anticoagulant tube containing 7 µL of 5 mM EDTA. For flow cytometry, 100 µL of peripheral blood was placed in 4 mL of ACK solution for 3 min at room temperature. In contrast, for cell sorting and sequencing, 1 mL of peripheral blood was placed in 5 mL of ACK solution for 5 min at room temperature, supplemented with 10 mL of PBS to inactivate ACK, followed by centrifugation and resuspended with 200 µL of FACS buffer.

### Flow cytometry

Cell suspensions were washed in PBS for all tissues, followed by live/dead staining (Zombie Aqua, Biolegend, 423102) at 1:500 dilution for 10 min at room temperature. Nonspecific antibody binding to cells was blocked by incubation with an anti-CD16/32 antibody at 4 °C for 15 min. Cells were stained by antibody cocktails at 4 °C for 25 min. Cells were analyzed on a BD LSRFortessa X20 (BD Biosciences) or a BD FACSCantoII (BD Biosciences). Flow data were further analyzed in FlowJo (10.8.1). For cell counting, 10 µL absolute counting beads (Invitrogen, C36950) were added to 300 µL cell suspension, and the absolute cell number was calculated according to the manufacturer’s protocol.

### Neutrophil isolation

Cell suspension from PB and Lung was first sorted via a neutrophil-negative selection kit (Miltenyi, 130-097-658) according to the manufacturer’s protocol. For bulk RNA sequencing, fluorescence-activated cell sorting was applied to all samples (BM, lung, PB) using BD Aria III FACS (BD Biosciences), and CD11b+ Ly6G+ cells were sorted for sequencing. Neutrophils were at a purity of > 99% for bulk-seq and a purity of 90% for single-cell sequencing. The dissociated single cells were stained with AO/PI for viability assessment using the Countstar Fluorescence Cell Analyzer.

### Neutrophil Depletion

For neutrophil depletion, we injected 100 µg of anti-mouse Ly6G antibody (BioXcell clone 1A8) intraperitoneally at 24 h or twice at 24 and 48 h prior to analysis, each time resulting in > 85% reduction in blood neutrophil counts compared to isotype controls (BioXcell clone 2A3).

### RNA Isolation and Library Preparation for Bulk RNA-seq

Total RNA was extracted using the TRIzol reagent (Invitrogen, CA, USA) according to the manufacturer’s protocol. RNA purity and quantification were evaluated using the NanoDrop 2000 spectrophotometer (Thermo Scientific, USA). RNA integrity (RIN) was evaluated using the Agilent 2100 bioanalyzer (Agilent Technologies, Santa Clara, CA, USA). Only samples with a RIN value > 7 were adopted to prepare the library. The libraries were constructed using the VAHTS Universal V6 RNA-seq Library Prep Kit according to the manufacturer’s instructions. The transcriptome sequencing and analysis were conducted by OE Biotech Co., Ltd. (Shanghai, China).

### Bulk RNA Sequencing Process and Analysis

The libraries were sequenced on an Illumina Novaseq 6000 platform, generating 150 bp paired-end reads. About 48,000,000 raw reads were generated for each sample. Raw reads of the fastq format were first processed using fastp, and the low-quality reads were removed to obtain clean reads. About 45,000,000 clean reads for each sample were retained for subsequent analyses. The clean reads were mapped to the reference genome using HISAT2. The FPKM of each gene was calculated, and the read counts of each gene were obtained by HTSeq count. The PCA analysis was performed using R (v 3.6.0) to evaluate the biological duplication of samples. Differential expression analysis was performed using DESeq2. Q value < 0.05 and log_2_(foldchange) > [1] was set as the threshold for significantly differential expression genes (DEGs). The GO analysis was performed according to DEGs using the R package clusterProfiler.

### Label-free proteomics analysis of neutrophils

One million isolated neutrophils for each sample were pelleted in 1.5-mL Eppendorf tubes and snap-frozen at −80°C. SDT (4% SDS, 100 mM Tris-HCl, pH 7.6) buffer was used for sample lysis and protein extraction. The amount of protein was quantified with the BCA Protein Assay Kit (Bio-Rad, USA). A volume of 20 µg of protein for each sample was mixed with 5 X loading buffer and boiled for 5 min. Subsequently, the proteins were separated on 4%–20% SDS-PAGE gel (constant voltage 180V, 45 min). Protein bands were visualized by Coomassie Blue R-250 staining. Protein digestion by trypsin was performed according to the filter-aided sample preparation (FASP) procedure described by Matthias Mann. The digest peptides of each sample were desalted in C18 cartridges, concentrated by vacuum centrifugation and reconstituted in 40 µL of 0.1% (v/v) formic acid.

LC-MS/MS analysis was performed on a Q Exactive mass spectrometer (Thermo Scientific) coupled to an Easy nLC (Proxeon Biosystems, now Thermo Fisher Scientific). The peptides were loaded onto a reverse phase trap column (Thermo Scientific Acclaim PepMap100, 100 μm*2 cm, nanoViper C18) connected to the reverse phase analytical column of C18 (Thermo Scientific Easy Column, 10 cm long, 75 μm inner diameter, 3 μm resin) in buffer A (0.1% formic acid in water) and separated with a linear gradient of buffer B (84% acetonitrile and 0.1% formic acid) at a flow rate of 300 nL/min. The mass spectrometer was operated in positive ion mode. MS data were acquired using a data-dependent top20 method, dynamically choosing the most abundant precursor ions from the survey scan (300–1800 m/z) for HCD fragmentation. The automatic gain control (AGC) target was set to 1e6, and the maximum injection time to 50 ms. The duration of dynamic exclusion was 30.0 s. Survey scans were acquired at a resolution of 60,000 at m/z 200, the resolution for HCD spectra was set to 15,000 at m/z 200, and the isolation width was 1.5 m/z. The normalized collision energy was 30 eV, and the underfill ratio, which specifies the minimum percentage of target value likely to be reached at maximum fill time, was defined as 0.1%. The instrument was run with the peptide recognition mode enabled.

MS raw data for each sample were combined and searched using MaxQuant 1.6.14 software for identification and quantification analysis. Cluster 3.0 (http://bonsai.hgc.jp/~mdehoon/software/cluster/software.htm) and Java Treeview software (http://jtreeview.sourceforge.net) were used to perform hierarchical clustering analysis. The Euclidean distance algorithm for similarity measure and average linkage clustering algorithm (clustering uses the centroids of observations) for clustering were selected when performing hierarchical clustering. A heat map was often presented as a visual aid in addition to the dendrogram. CELLO (http://cello.life.nctu.edu.tw/), a multi-class SVM classification system, was used to predict protein subcellular localization. To analyze the correlation between proteomics and transcriptomics data for lung and PB neutrophils, we compared proteomics data and RNA-sequencing data generated in this study. Consequently, we obtained a set of common proteins and genes from both datasets, compared proteomics log_2_FC and sequencing log_2_FC, and calculated the correlation statistics (Spearman rank-order correlation) statistics in R.

### Single-cell library construction and sequencing

The scRNA-Seq libraries were generated using the 10X Genomics Chromium Controller Instrument and Chromium Single Cell 3’ V3.1 Reagent Kits (10X Genomics, Pleasanton, CA). Briefly, cells were concentrated to approximately 1000 cells/µL and loaded into each channel to generate single-cell gel beads in emulsions (GEMs). After the RT step, the GEMs were broken, and barcoded cDNA was purified and amplified. The amplified barcoded cDNA was fragmented, A-tailed, ligated with adapters, and the index PCR amplified. The final libraries were quantified using the Qubit High Sensitivity DNA assay (Thermo Fisher Scientific), and the size distribution of libraries was determined using a High Sensitivity DNA chip on a Bioanalyzer 2200 (Agilent). All libraries were sequenced by the Illumina sequencer (Illumina, San Diego, CA) on a 150 bp paired-end run.

### scRNA-seq data processing and filtering

The quality of sequencing reads was evaluated using FastQC and MultiQC. Cell Ranger version 2.2.0 was used to align the sequencing reads (fastq) with the mouse mm10 mouse transcriptome and quantify the expression of transcripts in each cell. This pipeline resulted in a gene expression matrix for each sample, which records the number of UMIs for each gene associated with each cell barcode. The cellular specimens obtained from two distinct individuals were combined and subjected to combined analysis. Following the exclusion of low-quality cells and identification of neutrophils, a total of 20941 cells were obtained, with 9118 cells originating from peripheral blood and 11823 cells from lung tissue.

### Data integration

The single-cell sequencing data from lung and PB neutrophils were integrated using Canonical Correlation Analysis performed using the standard workflow from Seurat developers (https://satijalab.org/seurat/v3.2/integration.html). Integrated data were only used for principal component analysis (PCA), and all steps relied on PCA (clustering and UMAP visualization). All other analyses (for example, differential expression analysis) were based on the normalized data without integration.

### Unsupervised clustering

A total of 15 principal components were used to perform clustering and UMAP dimensional reduction. We performed the “FindClusters” function (resolution: 0.3) to cluster cells using the Louvain algorithm based on the same principal components as for the “RunUMAP” function.

### Clusters Identification

Herein, we used the “FindAllMarkers function (logfc.threshold= log[1.4]) based on normalized data to identify DEGs. Differential expression is determined based on the non-parametric Wilcoxon rank sum test. DEGs with adjusted P values > 0.05 were excluded. Subsequently, we identified each neutrophil subcluster according to their DEGs regarding former neutrophil nomination by Xie et al. and further confirmed their identification by correlation analysis.

### Scoring

We used the “AddModuleScore” function in Seurat to calculate the expression level of a set of genes within cells. Consequently, we assigned an IEG (Immediate Early Genes) expression score using the following genes as characteristics: Jun, Junb, Jund, Fos, Fosb, Atf3, Hsp90aa1, Hsp90ab1, Egr1, Hspa1a, Hspa1b, Hspa8, Zfp36, Cebpb, Cebpd, Socs3, Hspe1, Dusp1. Additional scores were assigned according to their GO accession.

### Electrophysiology

Whole-cell patch-clamp recordings were performed at room temperature using the Molecular Devices system (Axoclamp 200B, Digidata 1440, pCalmp 10). The membrane voltage was maintained at 60 mV. The pipette solution contained: 120 mM KCl, 30 mM NaCl, 1LmM MgCl_2_, 0.5LmM CaCl_2_, 5 mM EGTA, 4 mM Mg-ATP, and 10 mM HEPES, pH 7.4, osmolarity kept at 280–300LmOsm/L. The standard external solution (SS) contained: 150LmM NaCl, 5 mM KCl, 1LmM MgCl_2_, 2LmM CaCl_2_, 10 mM glucose, and 10LmM HEPES buffered to various pH values with Tris-base or HCl. The osmolarity of SS was kept at 300– 330LmOsm/L. Mechanical stimulation was applied to cells using a fire-polished glass pipette with a 3- to 5-µm tip diameter. Experiments were performed with dissociated single cells.

### Live cell imaging

Ratiometric imaging using Fluo-8 and CMRA dyes measured Ca^2+^ signals in wild-type and Vav1-Piezo1^-/-^ neutrophils. The selected bone marrow neutrophils were suspended with 2 µM CMRA (Invitrogen, C34551) and 5 µM Fluo-8 (Abcam 1345980-40-6) in HBSS (Solarbio, H1045) and seeded in a glass bottom cell culture plate coated with 0.0001% PDL (Beyotime, ST508) coated glass bottom cell culture dish (NEST, 801002), cultured at 37 5% CO_2_ for 30 min. The culture medium was changed to HBSS with calcium and magnesium before imaging. To stimulate a Piezo1-induced calcium influx, 30 µM Yoda1 (Sigma, SML1558) was topically added to the imaging field during live cell imaging. Olympus FLuoview FV3000 microscope equipped with a UplanSApo 20x/0.75 objective (Olympus). We acquired a time-lapse video to record the calcium influx every 3 s for 3 min at a single z.

### Immunofluorescence and confocal imaging

Lungs were infused with 600 µL UltraPure low melting point agarose (Invitrogen, 16520-050) (1.5% in PBS and prewarmed at 40 °C) through the trachea. Tissues were harvested and fixed overnight in 4% PFA (Servicebio, G1101) and sequentially dehydrated in 15% and 30% sucrose before embedding in the OCT compound (Sakura Finetek, 4583). The sections were cut to 30 µm on a Leica cryostat and permeabilized with 0.1% Triton X-100 for 2 h, then blocked with 0.1% Fc block and 2% BSA (Kingmorn, KS1090) for 1 h at room temperature. Antibodies were diluted in 0.2% Tween-20 in PBS, and the sections were stained at 4 °C in a dark, humidified chamber. Immune fluorescence antibodies can be found in this study in **Table S11**. After staining, the slides were mounted with an antifade mounting medium (Yeasen, 36307ES25) and examined on an Olympus FLuoview FV3000 microscope equipped with a UplanSApo 20x/0.75 objective (Olympus).

### Preparation of Bronchoalveolar Lavage Fluid (BALF)

Mice were anesthetized by intraperitoneal injection of 25 µL/g Avertin, then transcardially perfused with 10 mL cold PBS. The skin and muscles above the trachea were carefully removed. A small opening was cut in the trachea, and a syringe containing 1 mL of cold PBS was prepared with an indwelling needle. The indwelling needle was inserted into the small opening of trachea for about 1 cm. 1 mL PBS with 2.5 mM EDTA was slowly injected and then slowly withdrawn and collected to a tube, repeated three times. Cell suspensions were centrifuged and then resuspended in 100 μL FACS buffer for later use.

### RNA isolation, reverse transcription, and RT-PCR

Total RNA was isolated from sorted cells using TRIzol reagent (Invitrogen, 15596018). According to the manufacturer’s protocol, glycogen (Beyotime, D0812) was added as a co-precipitant reagent. According to the manufacturer’s recommendations, the cDNA was subsequently generated using a HiScript II Q RT SuperMix (Vazyme, R223). The pre-amplified cDNA was subjected to qPCR in which the amplified product was detected using AceQ Universal SYBR qPCR Master Mix (Vazyme, Q511-03) on a CFX384 Real-Time PCR Detection System (Bio-rad).

### Bacterial infection

The *Streptococcus pneumoniae* strain D39 was kindly provided by Dr. Jingren Zhang from Tsinghua University. The bacteria were cultured in Todd-Hewitt Broth (THB, Hopebio, HB0311-3) containing 0.5% yeast extract (Oxoid, LP0021) at 37 °C with 5% CO_2_ until reaching mid-logarithmic growth phase (OD600 of 0.6). The pulmonary infections were carried out by intratracheal (i.t.) injection, as described previously (An et al., 2022), with a final injection volume of 30 μL. Animal survival was monitored for seven days post-infection or until the humane endpoint. All infection experiments followed Shanghai Jiao Tong University School of Medicine guidelines.

### Statistical analysis and reproducibility

For most experiments, comparisons were made using a two-tailed, unpaired Student’s t-test. The values in each figure represent means ± SD. A *P*-value < 0.05 was considered statistically significant. All experiments were repeated at least three times. All statistical analyses and graphics were made using GraphPad Prism (GraphPad) and R (The R Project for Statistical Computing). ’n’ in the figure legends indicates the number of biologically independent replicates. The results of Western blot and immunofluorescent staining are representative of at least three biologically independent replicates. Unless specified, all outcomes were replicated independently on more than three separate times, producing similar results.

## Data availability

All sequencing data generated in this study have been deposited at NCBI’s Gene Expression Omnibus (GEO) repository. The published data used in this study were retrieved from the GEO (accession numbers GSE137540). The videos and other data supporting the findings of this study are available upon request from the corresponding author.

## Acknowledgments

We thank Core Facility of Shanghai Institute of Immunology, Shanghai Jiao Tong University School of Medicine for technical help with flow cytometry and intravital imaging. This work was supported by grants from The Ministry of Science and Technology of China (2020YFC2002800 to Jing Wang), the National Natural Science Foundation of China (81822020, 92042304, 31872737 to Jing Wang).

## Author Contributions

Jin W. performed most of the experiments, analyzed data and interpreted results. H.-X. K., W.-Y.Z., X.Q., W.-J.B., Y.-Y.W. and N.-J.M. helped with the experiments. A.T., D.-C.Z. and T.-L.X. helped with the manuscript. Jing W. designed and supervised the study. Jin W. and Jing W. wrote the manuscript.

