## supplementary figures for "Mechanical Force Imprints Neutrophils to Orchestrate Pulmonary Homeostasis"

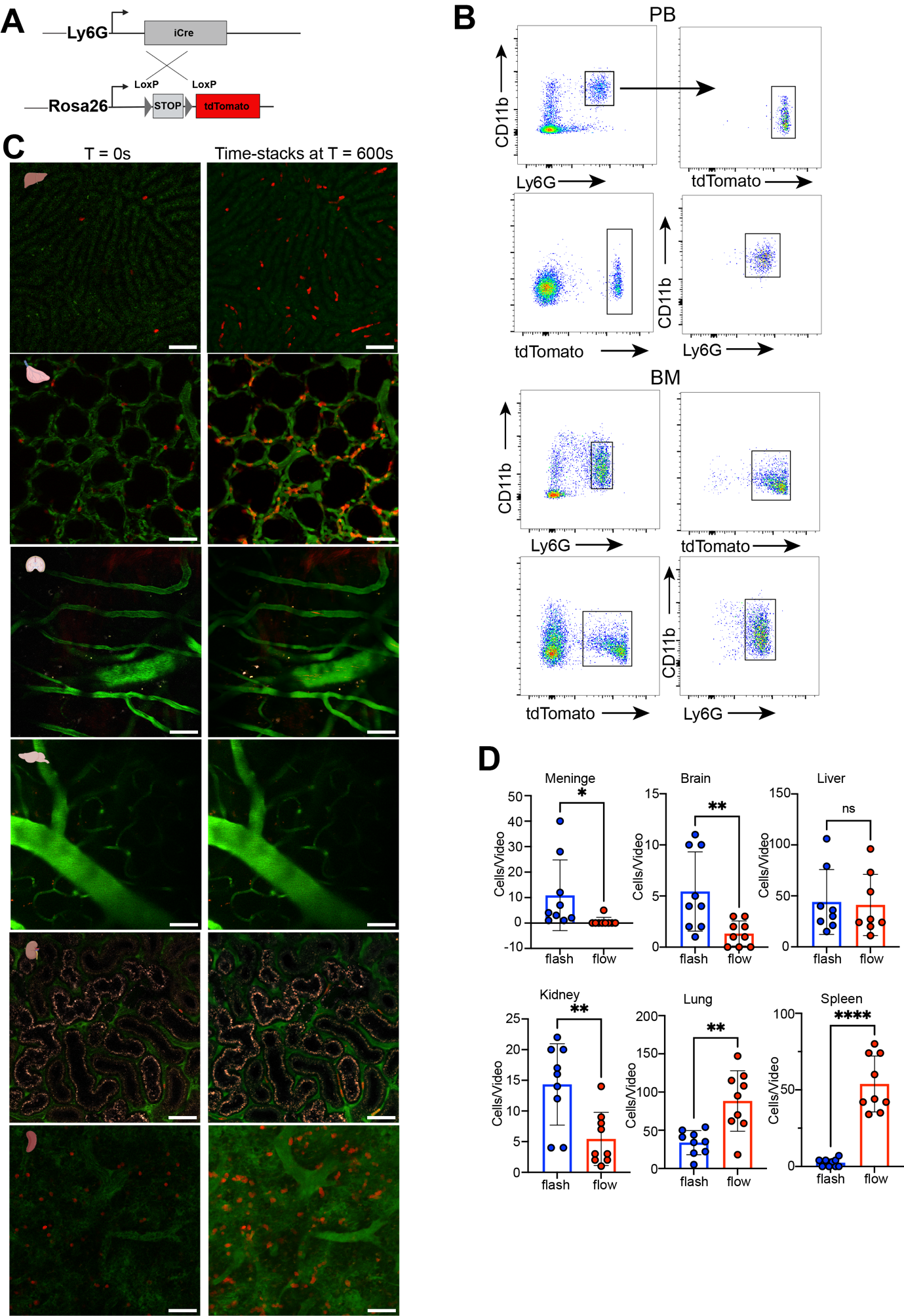

**Fig. S1. Intravital imaging analysis of neutrophil dynamics in different tissues.** (A) Construction strategy of Ly6G-tdTomato reporter mice. (B) Flow cytometry analysis of tdTomato expression in PB and BM neutrophils in Ly6G-tdTomato reporter mice. (C) Time-stacks from intravital imaging of different tissues in Ly6G-tdTomato reporter mice. Vasculature was labeled by i.v. injection of FITC-dextran 2000. (D) The number of neutrophils in different tissues visualized by intravital microscopy that display "flash" or "flow" migration behaviors.

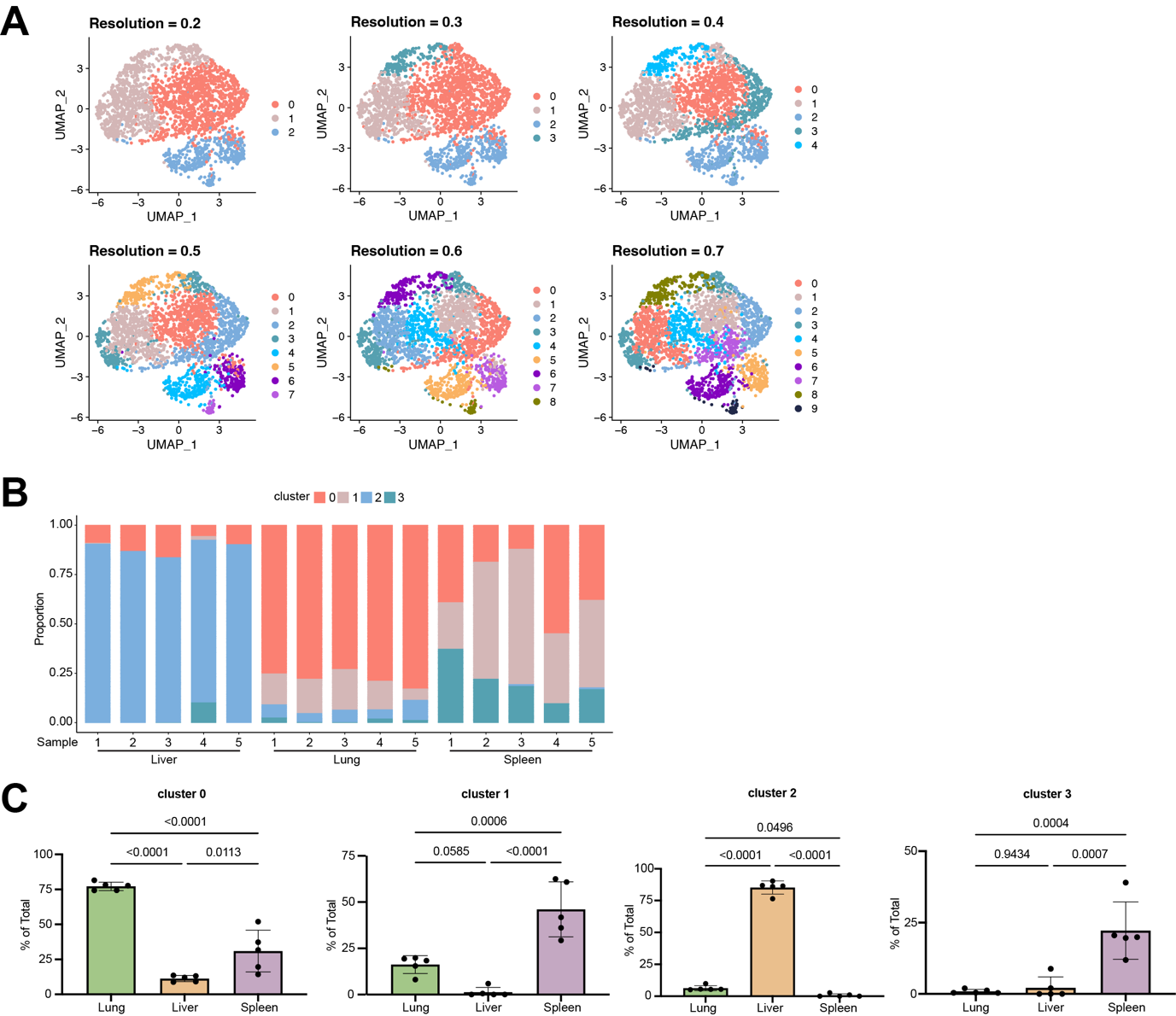
 **Fig. S2. Neutrophil clustering by behavior features.** (A) UMAP plots are generated using different resolution settings in the Suerat V4 R package. (B) Proportions of the four neutrophil clusters in different tissues across each individual mouse. (C) Percentage of different neutrophil subpopulations in different tissues. n = 5.

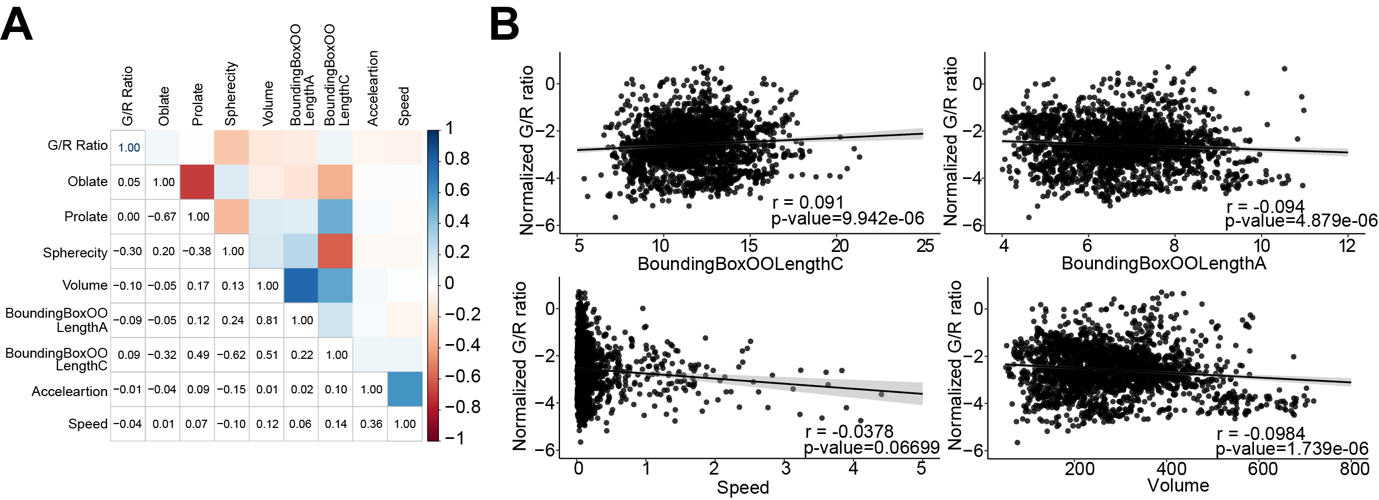
 **Fig. S3. Correlation analysis of neutrophil Ca^2+^ with other behavior features.** (A) Correlation matrix of paired variables assessed in the cellular behavior and Ca^2+^ analysis from intravital imaging experiments. *P* values are given, and the correlation coefficients are color coded. (B) Correlation between GCaMP6f/tdTomato intensity ratio (G/R ratio) and indicated parameters describing neutrophil kinetics and morphology traits in vivo.

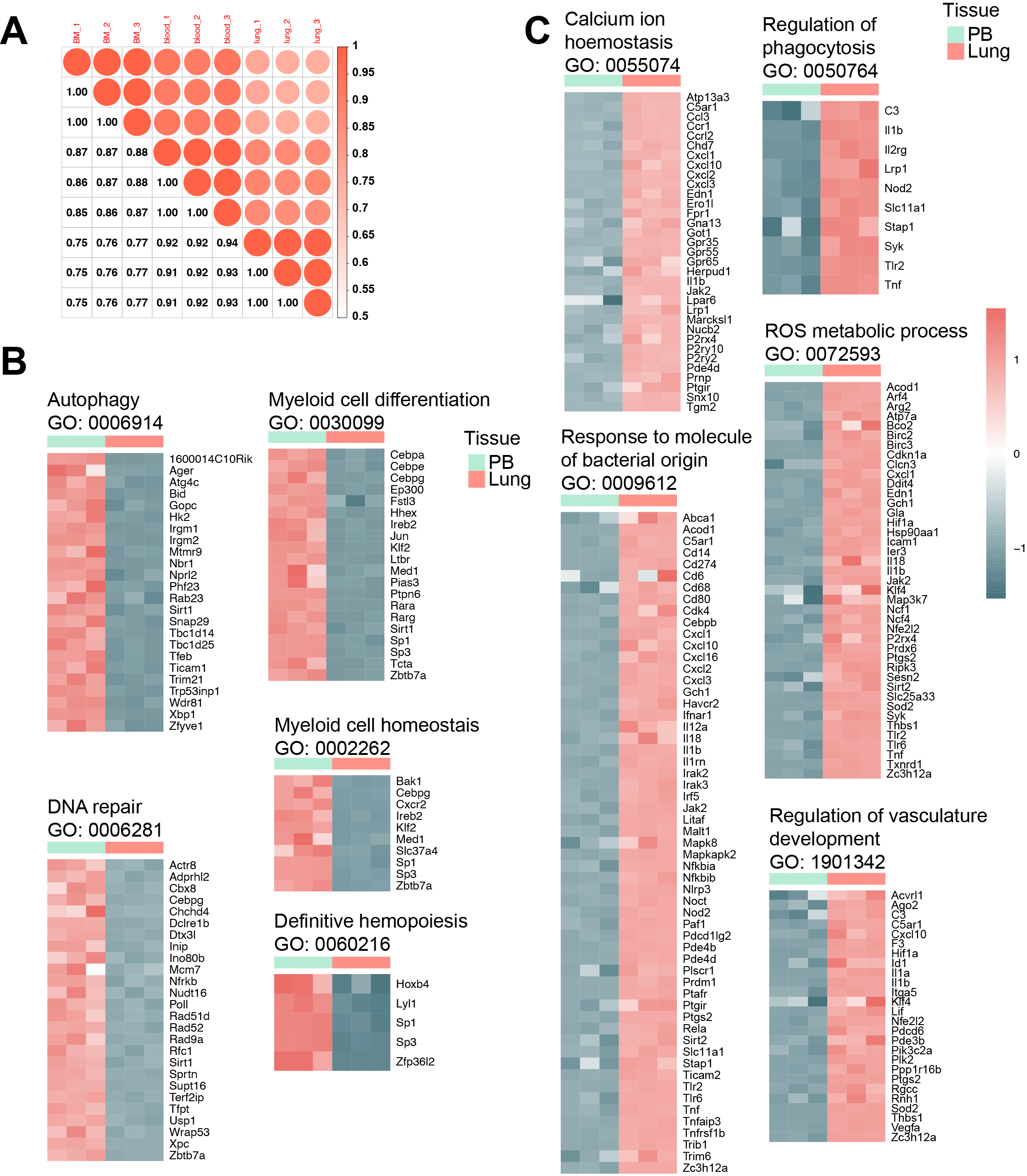

**Fig. S4. Pathways enriched in lung and blood neutrophils.** (A) Correlation matrix of gene expression levels between three biological replicates of each tissue. *P* values are given, and the correlation coefficients are color coded. (B–C) Examples of functionally defined gene subsets that are differentially expressed between the lung and peripheral blood neutrophils (adjust P value < 0.05). The log_2_ fold change to the geometric mean of FPKM + 1 is shown.

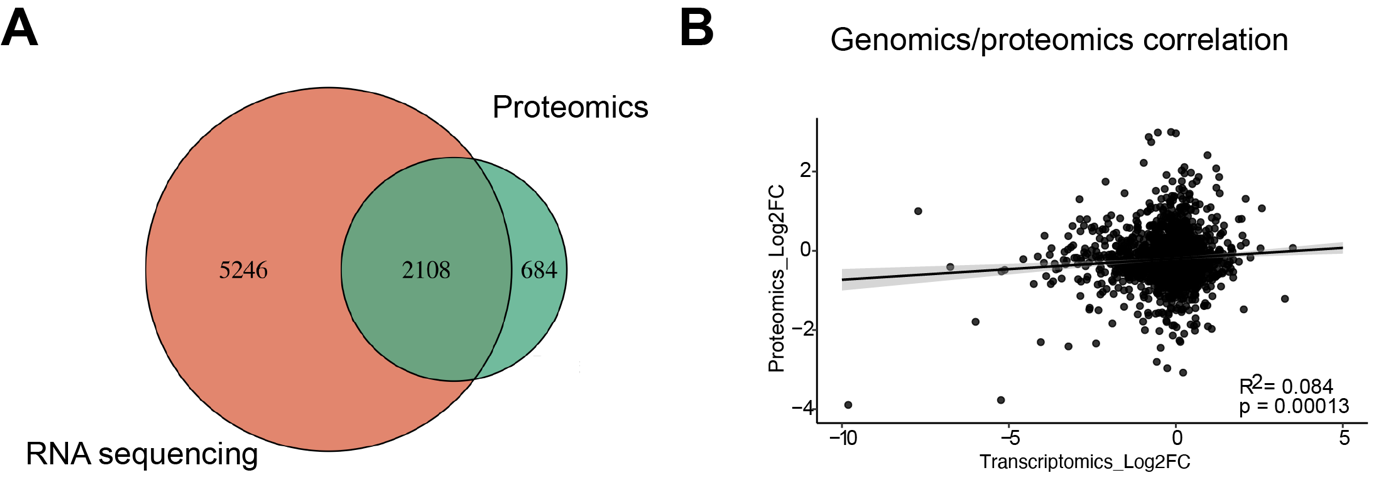

**Fig. S5. Analysis of the proteome of tissue-specific neutrophils.** (A) The Venn diagram represents unique and shared genes/proteins between transcriptome and proteome. (B) Correlation analysis (Spearman) of common proteins and genes change from paired proteomic and RNA sequencing analysis.

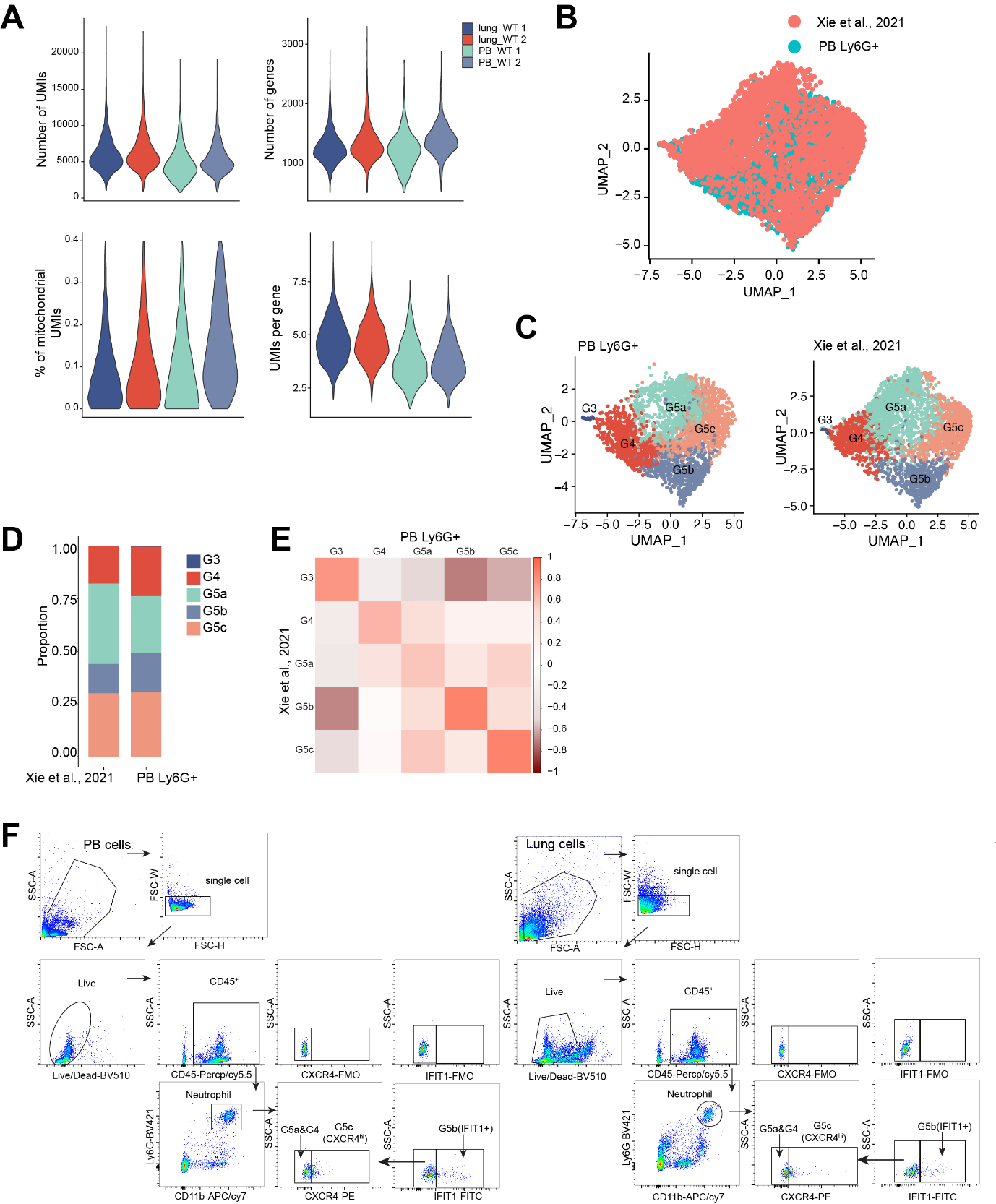

**Fig. S6. Comparison of scRNA-seq-defined neutrophil populations with previously reported neutrophil subpopulations.** (A) Violin plots of the number of UMIs, number of genes, mitochondria count percentage, and UMI per gene of all QC-passed cells in different organs and replicates. (B–C) Comparison of Ly6G+PM neutrophils population in our data with Gr1+ PB neutrophil populations in Dr. Hongbo Luo's data. UMAP of 9073 cells from our data and 3993 cells from Xie et al., 2021 colored by the data set (B) or cluster identity (C). (D) Neutrophil compositions in our data and Xie et al., 2021. (E) Correlation of scRNA-seq-defined neutrophil populations with the neutrophil subsets reported by Xie et al., 2021. Coefficient matrix showing deconvolution results of bulk profiles of indicated neutrophil subpopulations. Each column is normalized by column sums. (F) Flow cytometry analysis for G5a and G4 (IFIT1-CXCR4lo), G5b (IFIT1+), and G5c (IFIT1-CXCR4hi) neutrophils in PB (left panel) and lung (right panel).

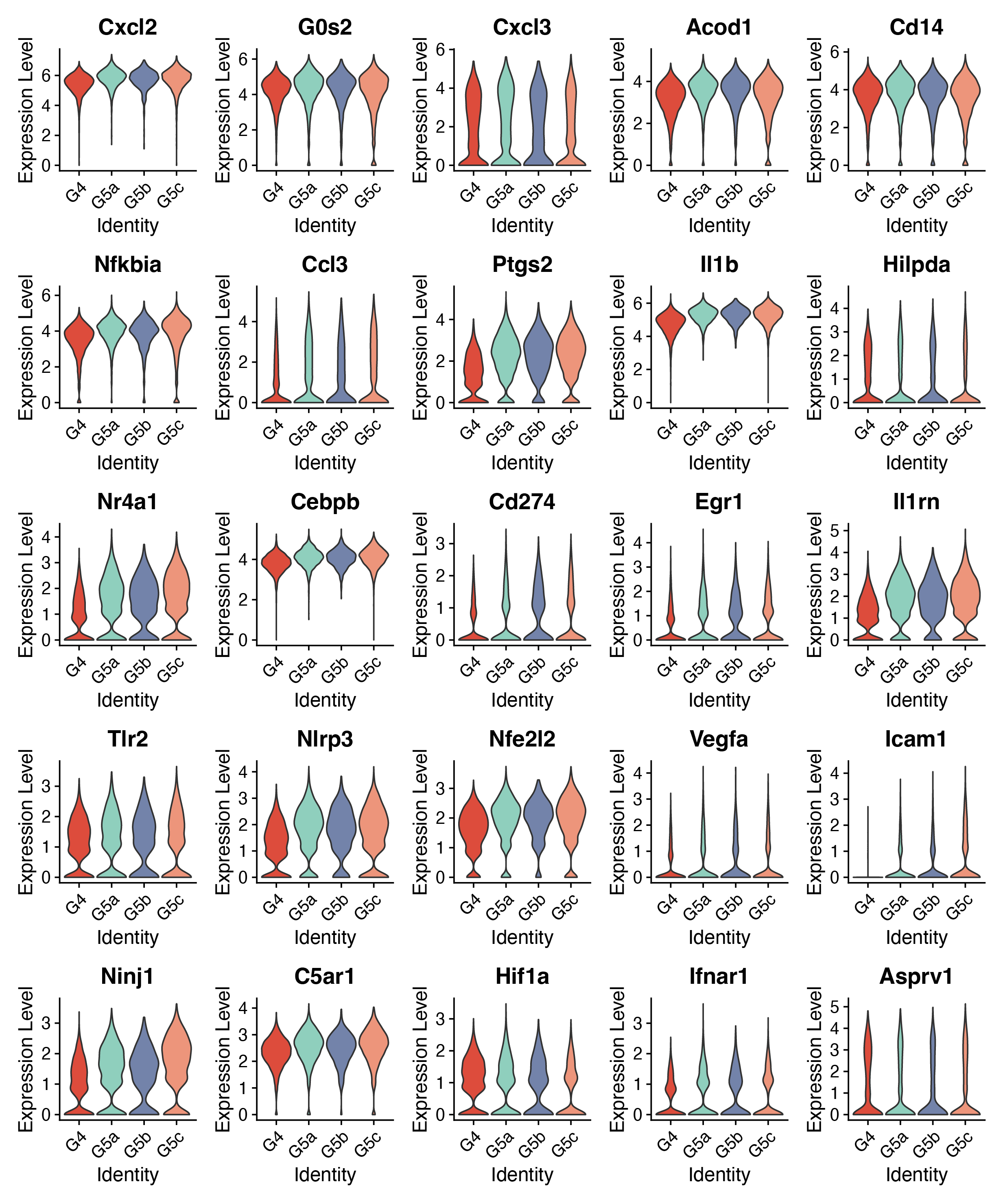

**Fig. S7. Expression of lung-specific signatures across neutrophil subpopulations.** Violin plots showing the expression levels of indicated genes in different neutrophil subsets.

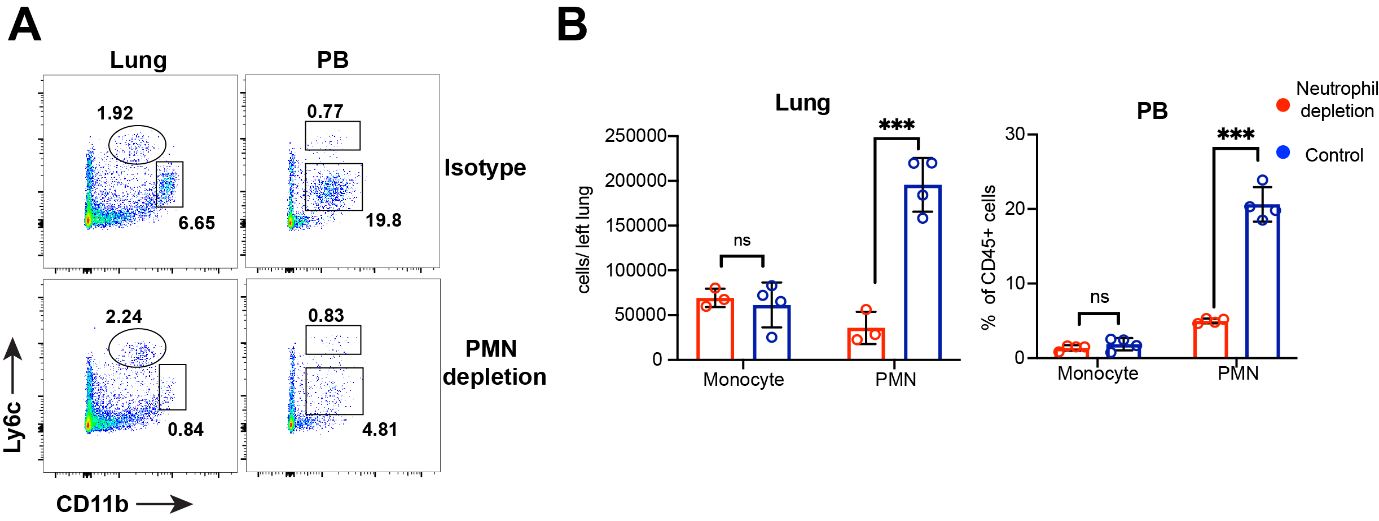

**Fig. S8. The efficiency of antibody-mediated neutrophil depletion.** (A) Representative flow cytometry analysis of neutrophils in lung and PB in isotype control (Isotype) and 1A8 antibody (PMN depletion) treated mice. (B) Absolute count (lung) or percentage (PB) of monocyte and neutrophil in isotype control-treated or 1A8 antibody-treated mice. n = 4.

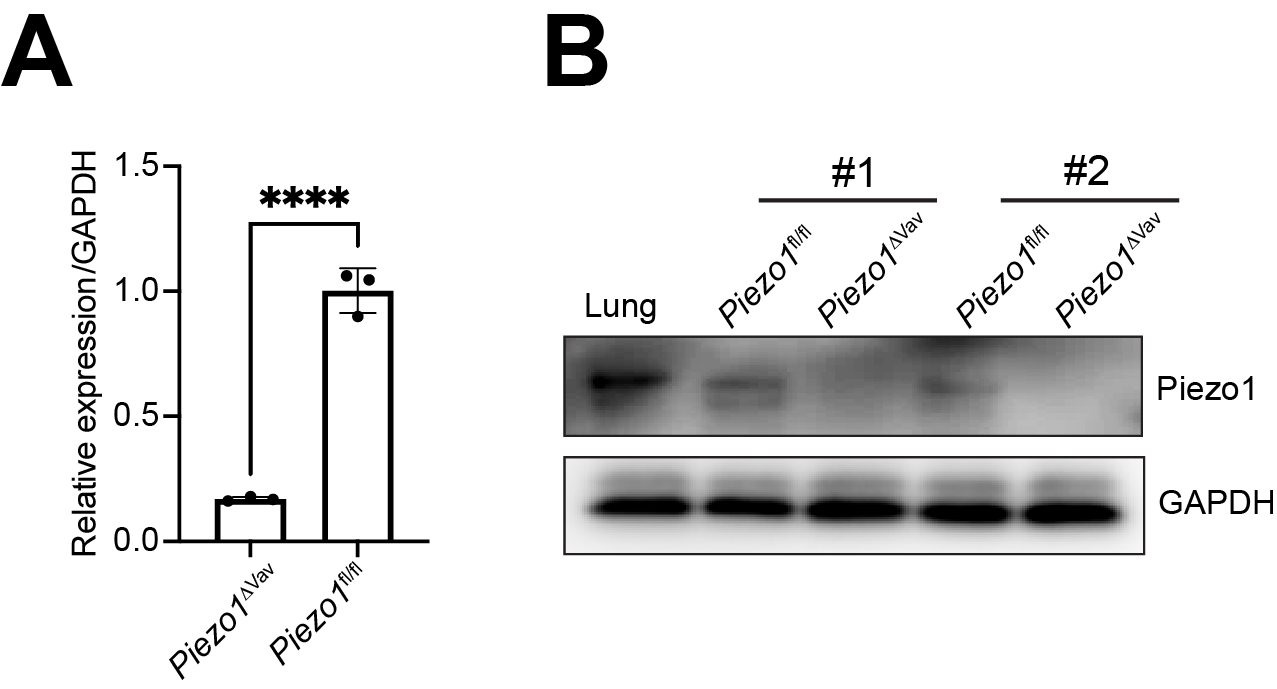

**Fig. S9. Verification of piezo1 deletion in neutrophils.** (A) RT-PCR confirmation of Piezo1 knockout in BM neutrophils from indicated mice. (B) Western blot of Piezo1 protein in BM neutrophils from indicated samples. Two pairs of littermates were analyzed.

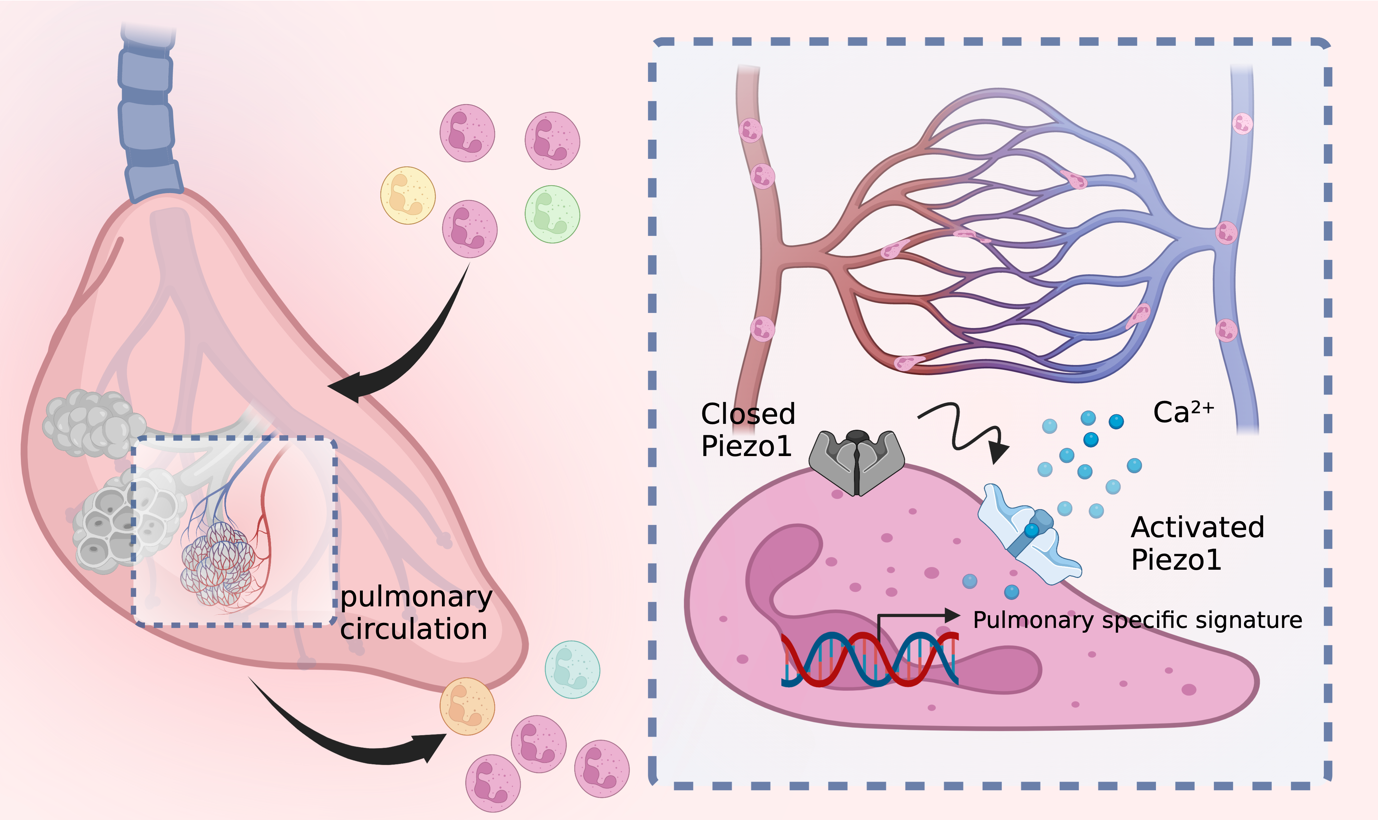

**Fig. S10.** Graphical abstract. When migrating across the capillary network in the lung, due to the size mismatch, neutrophils have to deform to pass through the narrow blood vessels. The Mechanical tension generated during this progress can be sensed by Piezo1, which allowed the influx of Ca^2+^ and altered the downstream gene expression. This imprints neutrophils to express lung specific signatures. As a consequence, neutrophils display lung-specific functions such as promoting angiogenesis. Lacking of PIEZO1 on neutrophils results in impaired capillary angiogenesis.

**MOV. S1. Intravital imaging of neutrophils in different tissues.** Intravital imaging was performed in indicated tissues in Ly6G^tdTom^ mice. Mice were treated intravenously with FITC-dextran to label blood vessels. Representative of five independent mice per organ.

**MOV. S2. Neutrophils display Ca^2+^ transient while migrating across the lung.** Intravital imaging of lungs in Ly6G^Salsa6f^ mice. Mice were treated intravenously with FITC-dextran (grey) to label blood vessels. Arrows indicate cells that display transient Ca2+. Representative of five independent mice.

**MOV. S3. No Ca^2+^ events in neutrophils in the spleen and liver.** Intravital imaging of spleen and liver in Ly6G^Salsa6f^ mice. Representative of five independent mice.

**MOV. S4. Neutrophils display prolonged Ca^2+^ signaling in response to infection.** Intravital imaging of lungs in Ly6G^Salsa6f^ mice 30 min after i.t. treatment of *Streptococcal pneumonia*. Representative of five independent mice.

**MOV. S5. Comparison of pulmonary neutrophils migration behaviors and Ca^2+^ signaling in *Salsa6f*^ΔVav1^ (WT) and *Salsa6f*-*Piezo1*^ΔVav1^ (Piezo1-cKO) mice.** Intravital imaging of lungs in *Salsa6f*^ΔVav1^ and *Salsa6f*-*Piezo1*^ΔVav1^ mice. Mice were treated intravenously with fluorescent anti-Ly6G to label neutrophils. Representative of five independent mice.
